# The ChlorON Series: Turn-on Fluorescent Protein Sensors for Imaging Labile Chloride in Living Cells

**DOI:** 10.1101/2022.08.13.503807

**Authors:** Jasmine N. Tutol, Whitney S. Y. Ong, Shelby M. Phelps, Weicheng Peng, Helen Goenawan, Sheel C. Dodani

## Abstract

Beyond its role as the queen electrolyte, chloride can also serve as an allosteric regulator or even a signaling ion. To illuminate this essential anion across such a spectrum of biological processes, researchers have relied on fluorescence imaging with genetically encoded sensors. In large part, these have been derived from the green fluorescent protein found in the jellyfish *Aequorea victoria*. However, a standalone sensor with a turn-on intensiometric response at physiological pH has yet to be reported. Here, we address this technology gap by building on our discovery of mNeonGreen (mNG) derived from lanYFP found in the cephalochordate *Branchiostoma lanceolatum*. Targeted engineering of two non-coordinating residues in the chloride binding pocket of mNG unlocks the ChlorON series. *In vitro* spectroscopy reveals that the binding of chloride tunes the chromophore environment to give rise to the turn-on response. We further showcase how this unique sensing mechanism can be exploited for directly imaging labile chloride in living cells with spatial and temporal resolution, accelerating the path forward for fundamental and translational aspects of chloride biology.

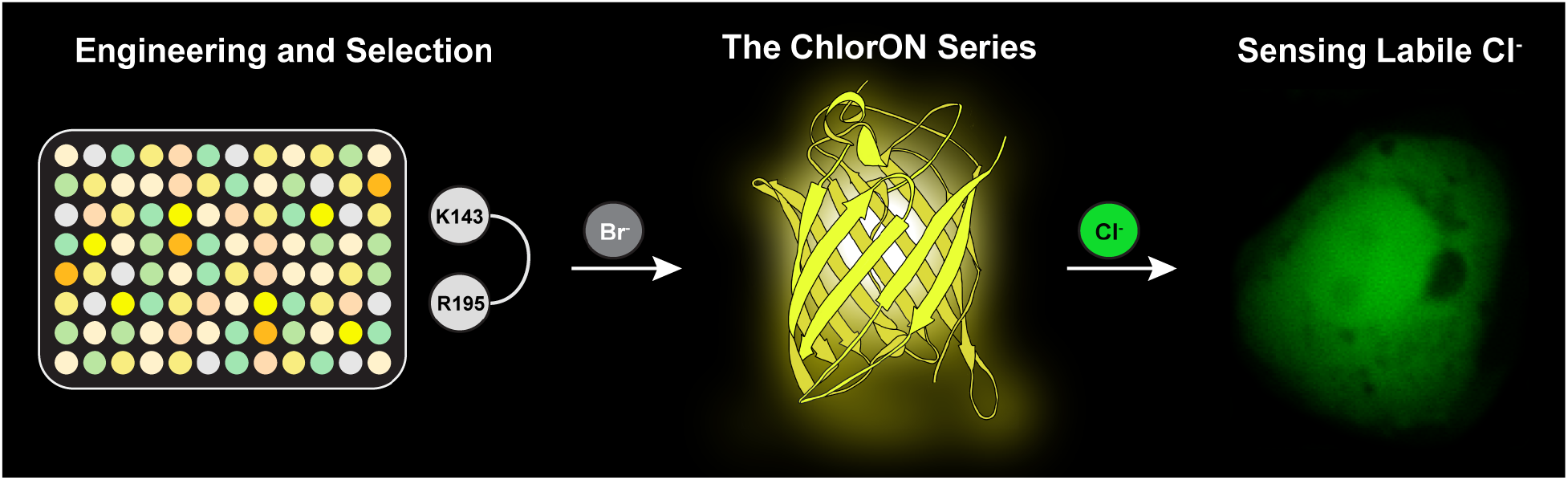

## Introduction

Inarguably, the chloride (Cl^-^) anion is an essential nutrient in physiology.^1^ Membrane-bound transporting proteins are resident on virtually every cell type, tuning chloride over a wide concentration range (3–110 mM) in extracellular, intracellular, and subcellular spaces.^2–9^ Given this redundancy, the activity of chloride channels, exchangers, and transporters can be passive or controlled by a myriad of stimuli, including cations (e.g., Na^+^, Ca^2+^), ligands (e.g., GABA, cAMP), or post-translational modifications.^3,4,7,10–13^ In turn, chloride is linked to biological functions such as circadian rhythms, electrolyte balance, fluid secretion, and innate immunity.^1,7,14–16^ Indeed, disruptions in the activity, expression, and localization of chloride transporting proteins can lead to dysregulated chloride levels in disease states ranging from cancers, cardiac dysfunction, cystic fibrosis, kidney stones, and neurological disorders.^17–24^ Across these contexts, the canonical view of chloride as a counterion is being re-shaped. Recent evidence ascribes functional significance to chloride as a signaling ion that can allosterically regulate protein function and even control gene expression.^11,12,14,25–27^ Thus, indicating that cells can maintain and mobilize chloride from distinct pools.

Against this backdrop, we are engineering a modern arsenal of technologies to investigate chloride in diverse cellular processes. Historically, chemical precipitation assays, electrophysiology, and radiolabeling alongside molecular blockers and ionophores have provided a readout of chloride transport.^2,28–35^ To gain such insights in large populations of cells, Illsley and Verkman developed the first fluorescent sensors to image chloride more than thirty years ago.^28,36^ Ion-pairing between a quinolinium or an acridinium fluorophore with halides results in collisional fluorescence quenching.^9,28,36–43^ Despite technical limitations, these sensors have stood the test of time with recent developments addressing subcellular targeting.^6,28,36,41,44–48^

Similarly, genetically encoded biosensors for chloride have been derived from the green fluorescent protein (GFP) found in the jellyfish *Aequorea victoria* (avGFP).^49–51^ Wachter and Remington first uncovered five mutations – S65G, V68L, S72A, H148Q, and T203Y – in and around the tripeptide chromophore that created an anion binding pocket and conferred sensitivity in the resulting avYFP-H148Q variant (Figure S1).^52–54^ Chloride as well as less abundant biological anions (e.g., Br^-^, I^-^, NO_3_^-^) shift the chromophore equilibrium or p*K*_a_ in a dose-dependent manner away from the fluorescent phenolate form to the non-fluorescent phenol form, generating a turn-off intensiometric response at physiological pH.^53–55^ This sensing mechanism enabled the first imaging application of a genetically encoded sensor for chloride in living cells.^55^ From these landmark studies, the avGFP sequence has been diversified through mutagenesis resulting in a palette of intensiometric and ratiometric biosensors.^5,9,56–58^ Included are avYFP-H148Q variants, E^2^GFP, Cl-Sensor, SuperClomelon, and LSSmClopHensor, to name a few (Figure S1).^54,59–70^ Over the last five years, we and others have explored the rich biodiversity of the GFP family for starting points with unique features that can be optimized for cellular applications through protein engineering.^71–74^

In 2019, we reported the discovery of mNeonGreen (mNG) from the yellow fluorescent protein (lanYFP) found in the cephalochordate *Branchiostoma lanceolatum* as a turn-on intensiometric sensor for chloride.^72,75^ Unlike the aforementioned sensors, chloride shifts the chromophore equilibrium from the non-fluorescent phenol form to the fluorescent phenolate form (Figure 1A). We further demonstrated that this sensing mechanism is linked to the substitution at position 195 above the chromophore, which is homologous to the key tyrosine at position 203 in the avGFP-based sensors.^53,64,72^ However, the turn-on response is only apparent at acidic pH. This can be rationalized based on the X-ray crystal structures of mNG (Figure 1B). At acidic pH, chloride (70% occupancy) interacts directly with residues H62, R88, S153, T173 and Y175, and indirectly through water with residues K143 and R195 (Figure 1B). On the contrary, at basic pH, chloride is largely absent (30% occupancy) and can be replaced with water as the K143 residue is modified with a carboxyl group, introducing a negative charge (Figure 1C).^75,76^ It is important to point out that there is no evidence for this lysine carboxylation in solution, likely given its transient and reversible nature.^77^ Nonetheless, all these observations suggest that position 143 in the context of position 195 could play a key role in chloride coordination and sensing at physiological pH. In this study, we explore this hypothesis and unlock unprecedented function in the ChlorON series – the first turn-on intensiometric sensors for directly imaging labile chloride with spatial and temporal resolution in living cells.

**Figure 1.**
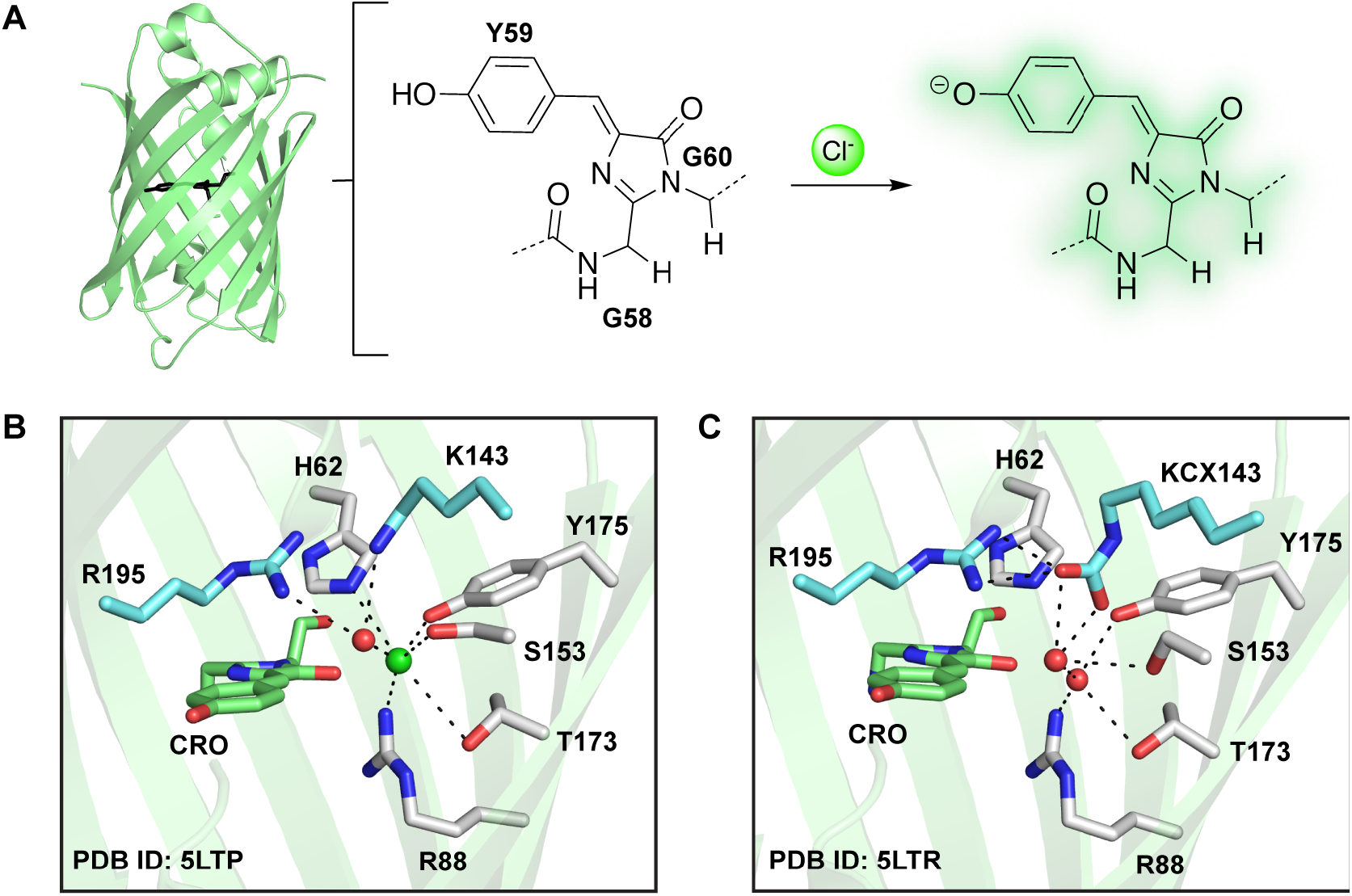
(A) The overall structure of mNeonGreen (mNG) bound to chloride (green sphere) (PDB ID: 5LTP) with the chromophore shown as black sticks is shown on the left. The chloride-dependent conversion of the chromophore from the non-fluorescent phenol to the fluorescent phenolate form, generating the turn-on response at pH 4.5 is shown on the right. (B) Comparison of the chloride binding pockets in mNG crystallized (B) at pH 4.5 and (C) pH 8.0. For each structure, the residues that make up the chromophore (green) and residues within 4 Å of the chloride ion (green sphere) with waters (red sphere) are shown. All residues are labeled with the single amino acid abbreviation and corresponding position number. The hydrogen bonding and electrostatic interactions are shown as dashed lines. Abbreviations: KCX, carboxylated lysine.

## Results

### Engineering mNG into the ChlorONs

To explore the effect of mutations at the K143 and R195 sites in the mNG parent, we employed a double site-saturation mutagenesis strategy using the 22-c trick to sample all possible amino acid substitutions and combinations (Figure 2 and S2). Compared to randomization approaches with NNK/S degenerate codons, the 22-c trick reduces the codon degeneracy and thus, the sampling size by at least half.^78^ Two sets of primers with the NDT/VHG/TGG codes for each targeted site were designed and used to clone the library in a single polymerase chain reaction (Table S1 and S2). The resulting library was transformed into *Escherichia coli* and 1,880 unique clones (>95% coverage) were randomly selected and expressed in a 96-well plate format for screening (Figure 2). To unlock the anion-sensitivity of mNG at physiological pH, we explored an anion walking strategy by first measuring the fluorescence response of the variants to bromide at pH 8 using an enzyme-based lysis with lysozyme and then to chloride at pH 7 and 8 using a sonication-based lysis. The rationale for our approach was motivated by the concept of substrate walking traditionally used in biocatalysis.^79^ In comparison to chloride, bromide has a larger ionic radius (Cl^-^ 1.8 Å < Br^-^ 2 Å) and lower dehydration enthalpy (Cl^-^ 365 kJ/mol > Br^-^ 335 kJ/mol).^80,81^ These physical properties typically result in higher affinity complexes. Indeed, this is observed with mNG at pH 4.5 (*K*_d_ = 1.8 ± 0.2 mM for Br^-^ versus *K*_d_ = 9.8 ± 0.3 mM for Cl^-^).^72^ By relying on anion walking, we identified six variants with a turn-on fluorescence response to both bromide and chloride in cell lysate: K143Y/R195I, K143R/R195L, K143R/R195I, K143W/R195L, K143G/R195K, and K143L/R195I (Figure S3, Table S3). Based on these data, the top three variants – K143W/R195L (ChlorON-1), K143R/R195I (ChlorON-2), and K143R/R195L (ChlorON-3) – were selected for further *in vitro* characterization (Table S3).

**Figure 2.**
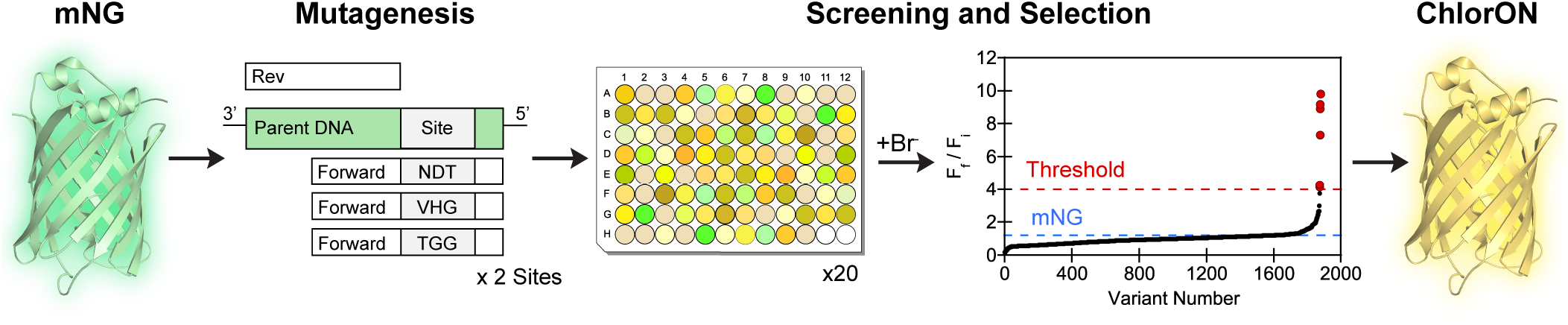
Protein engineering workflow to generate the ChlorON sensors. Double-site saturation mutagenesis was carried out at K143 and R195 in the mNG parent followed by transformation, expression, and fluorescent plate reader screening of the library in *E. coli* lysate in the absence and presence of 100 mM NaBr in 25 mM sodium phosphate buffer at pH 8. Variants with at least a ∼4-fold turn-on fluorescence response were selected for further characterization with 100 mM NaCl in 25 mM sodium phosphate buffer at pH 7 and 8. The screening results are summarized in Figure S3 and Table S3.

### Characterization of the ChlorONs

To determine if the results from the cell lysate screening translate in purified form, we collected the absorption and emission spectra of the ChlorONs at pH 7. Each sensor was expressed in *E. coli* and purified on a preparative scale using affinity and size-exclusion chromatography. As expected, the mutations introduced at positions 143 and 195 in the binding pocket did not affect the monomeric oligomerization state (Figure S4). All three sensors have broad absorption maxima centered at ∼480 nm. In the presence of 197 mM NaCl, the relative intensity of the absorption maximum remains unchanged for ChlorON-1 but decreases and shifts to 505 nm for ChlorON-2 and ChlorON-3 (Figure S6–S8, Table S4). Both absorption maxima correspond to the phenolate form arising from two different vibrational modes as observed with mNG and other GFPs.^82–85^ Upon excitation of the phenolate form at 485 nm, each sensor has a quenched emission maximum centered at 515 nm (Figure 3, Table S4). Titration with up to 197 mM NaCl does not shift the emission maxima but triggers a turn-on fluorescence response that can be ranked as follows: ChlorON-1 (45-fold) > ChlorON-2 (27-fold) > ChlorON-3 (20-fold). Indeed, the brightness of each sensor increases and tracks with the relative chloride binding affinities (*K*_d_): ChlorON-1 (0.65, *K*_d_ = 285 ± 59 mM) < ChlorON-2 (1.9, *K*_d_ = 55 ± 5 mM) < ChlorON-3 (2.8, *K*_d_ = 30 ± 1 mM) (Figure 3, Table S4). Under similar testing conditions, no response is observed with the mNG parent (Figure S5).

**Figure 3.**
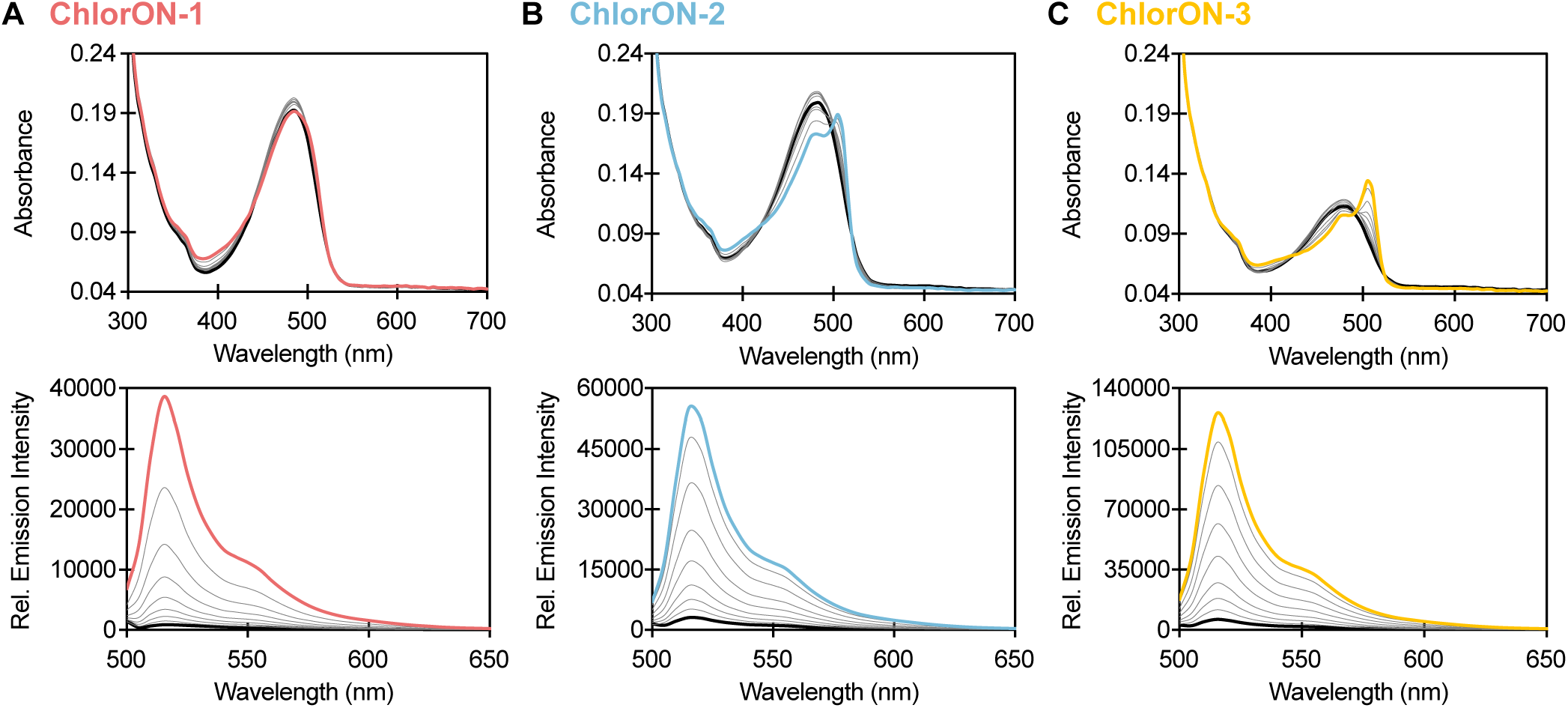
Spectroscopic characterization of the ChlorON sensors. Absorption (top row) and emission spectra (bottom row) of 10 μM (A) ChlorON-1, (B) ChlorON-2 and (C) ChlorON-3 in the presence of 1 (bold black), 2, 3.9, 6.9, 13.3, 25.5, 50, 99, and 197 mM (red, blue, and yellow, respectively) NaCl in 50 mM sodium phosphate buffer at pH 7 (λ_ex_ = 485 nm, λ_em_ = 500–650 nm). The average of four technical replicates from two protein preparations is shown. Plots with the average and standard error of the mean are reported in the Supporting Information (Figure S6–S8, Table S4).

### Characterization of the ChlorONs as a function of pH

To understand how the unique sensing mechanism of the ChlorONs could be connected to pH, we next characterized the spectroscopic properties of each sensor from pH 3.5 to 8.5 in the absence and presence of 197 mM NaCl (Figure 4). All three sensors have pH-dependent absorption spectra that reflect the chromophore equilibrium (Figure S10–S12). Interestingly, excitation of the phenol form (λ_ex_ = 400 nm) results in broad emission spectra with two maxima corresponding to the phenol (λ_em_ = 465 nm) and phenolate (λ_em_ = 515 nm) forms. This observation indicates that the local chromophore environment could stabilize an emissive phenol form and promote excited state proton transfer to generate the phenolate form.^86,87^ However, excitation of the phenol form was not investigated in greater detail given the low signal-to-noise ratios. Excitation of the phenolate form results in a single emission maximum at 515 nm that does not shift as a function of pH nor chloride but rather intensity for all three sensors (Figure S10– S12). These data were fitted to the Henderson-Hasselbalch equation to determine the chromophore p*K*_a_s. Notably, the emission intensity of ChlorON-1 does not vary to a large extent with increasing pH (p*K*_a_ could not be determined or ND) but increases with the binding of chloride from pH 5.5 to 9 (p*K*_*a*_ *=* 6.0 ± 0.1). Similarly, ChlorON-2 and ChlorON-3 respond to chloride across this pH range, but the emission intensity profiles are bell-shaped, giving rise to two p*K*_a_s. In the presence of chloride, the chromophore p*K*_*a*_s increase from 4.6 ± 0.1 and 6.1 ± 0.1 to 5.7 ± 0.1 and 7.6 ± 0.1 for ChlorON-2; from 4.8 ± 0.2 and 6.1 ± 0.2 to 6.0 ± 0.01 and 7.9 ± 0.1 for ChlorON-3 (Figure S11 and S12, Table S4).

**Figure 4.**
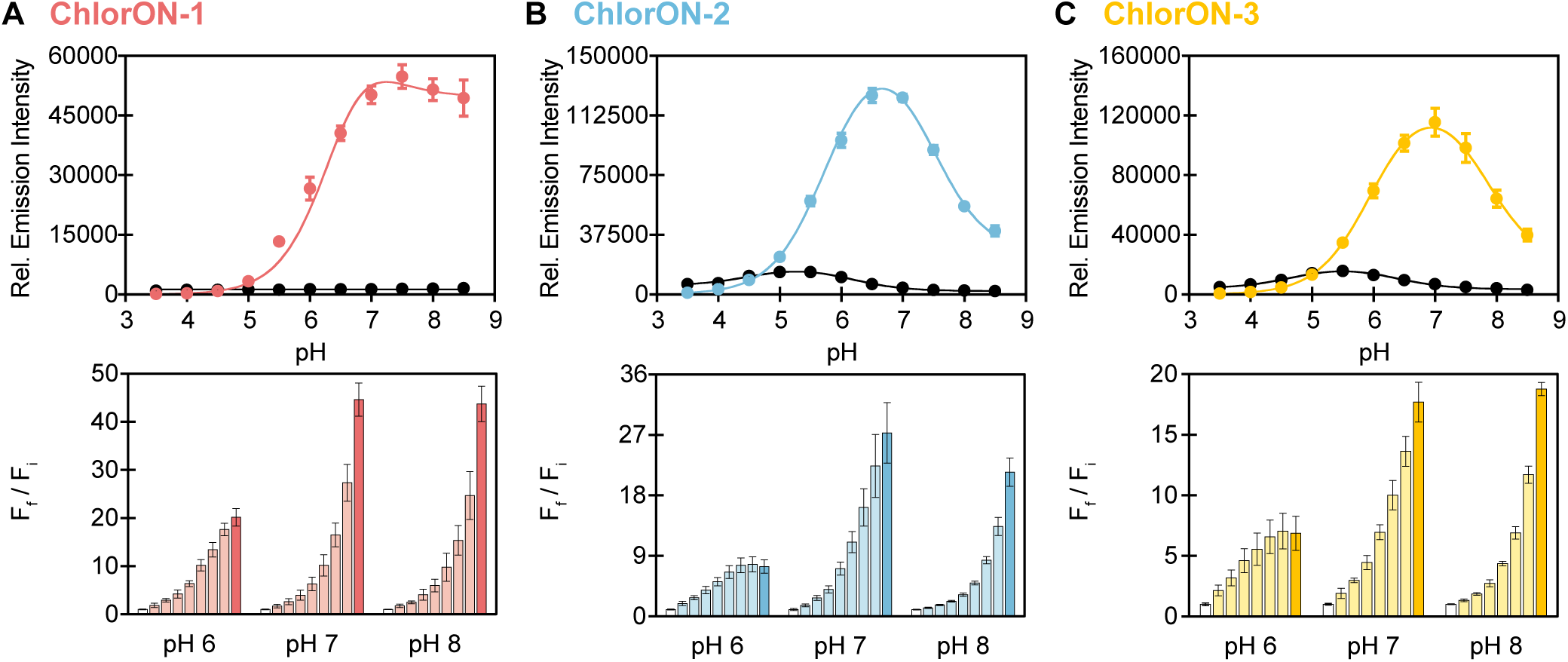
The pH profiles of the ChlorON sensors. The pH-dependent response of (A) 10 μM ChlorON-1 (left), ChlorON-2 (middle), and ChlorON-3 (right) to 1 mM (black circles) and 197 mM (red, blue, or yellow circles) chloride in 50 mM sodium acetate buffer from pH 3.5–5.5 and 50 mM sodium phosphate buffer from 5.5–8.5 (λ_ex_ = 485 nm, λ_em_ = 515 nm). The data is fitted to the Henderson-Hasselbach equation (black curves) to determine the p*K*_a_ values. (B) The fluorescence response of 10 μM ChlorON-1 (left), ChlorON-2 (middle), and ChlorON-3 (right) to 1 (white bar, F_i_), 2, 3.9, 6.9, 13.3, 25.5, 50, 99, and 197 mM (red, blue, or yellow bars) NaCl (F_f_) in 50 mM sodium phosphate buffer at pH 6, 7, and 8 (λ_ex_ = 485 nm, λ_em_ = 515 nm). The average of four technical replicates from two protein preparations with standard error of the mean is reported (Figure S6–S8, S10–S12, Table S4).

Based on this, we further characterized the ChlorONs with up to 197 mM NaCl at pH 6 and pH 8. Under these constant pH conditions, the absorption spectra indicate that each sensor is in the phenolate form at 480 nm. Chloride can tune the chromophore to the phenolate and phenol forms at 480 nm (ChlorON-1) and 505 nm (ChlorON-2 and ChlorON-3) and 400 nm, respectively. This effect is more pronounced at pH 6, in line with the calculated p*K*_a_s. Despite the complex nature of the absorption profiles, all three sensors have a turn-on fluorescence response to chloride and can be ranked as follows at pH 6: ChlorON-1 (20-fold) > ChlorON-2 (7.4-fold) > ChlorON-3 (6.2-fold); at pH 8: ChlorON-1 (44-fold) > ChlorON-3 (29-fold) > ChlorON-2 (21-fold) (Figure 4, Table S4). One noteworthy observation is that with increasing pH each sensor has a larger dynamic range while the chloride binding affinity decreases: ChlorON-1 (*K*_d_ = 39 ± 5 at pH 6; *K*_d_ = ND at pH 8), ChlorON-2 (*K*_d_ = 7.5 ± 1 mM at pH 6; *K*_d_ = 228 ± 25 mM at pH 8), and ChlorON-3 (*K*_d_ = 4.4 ± 0.9 mM at pH 6; *K*_d_ = 169 ± 79 mM at pH 8) (Table S4).

### Characterization of the anion selectivity for the ChlorONs

Since the selection of the ChlorONs relied on the anion coordination plasticity of the mNG parent, we characterized the response of all three sensors to a range of halides and oxyanions at pH 7.^72,88^ Concentration-dependent spectral changes are observed in the presence of bromide, iodide, nitrate, and sulfate, but not acetate, citrate, and phosphate (Figure S13–S19). Thus, indicating that anions larger than chloride with a particular shape and charge can bind to the same pocket and affect the chromophore environment, albeit to different extents. As described above for chloride, more pronounced absorption changes are observed for ChlorON-2 and ChlorON-3 than ChlorON-1. Specifically, bromide, iodide, and nitrate tune the chromophore equilibrium from the phenolate form at 480 nm to the phenolate form at 505 nm, and interestingly, to the phenol form at 400 nm. However, the addition of sulfate gives rise only to the latter.

To compare the fluorescence response of the sensors to each anion, excitation was only provided at 485 nm. As can be seen in Figure 5, ChlorON-1 is more selective for chloride (45-fold, *K*_d_ = 285 ± 59 mM) and bromide (26-fold, *K*_d_ = 204 ± 38 mM) than iodide (2.6-fold, *K*_d_ = ND), nitrate (2.2-fold, *K*_d_ = ND), and sulfate (ND, *K*_d_ = ND) (Figure S6, S16–S19, Table S5). Comparatively, ChlorON-2 and ChlorON-3 have measurable fluorescence responses with relatively higher binding affinities for each anion. Notably, fluorescence quenching is only observed in the presence of sulfate, which is in line with the absorption changes described above. The fluorescence response can be ranked as follows for ChlorON-2: bromide (30-fold, *K*_d_ = 43 ± 0.6 mM) ≈ chloride (27-fold, *K*_d_ = 55 ± 5 mM) > nitrate (9.1-fold, *K*_d_ = 117 ± 18 mM) > iodide (6.3-fold, *K*_d_ = 86 ± 4 mM) > sulfate (68%, *K*_d_ = 17 ± 2 mM); for ChlorON-3: chloride (20-fold, *K*_d_ = 30 ± 1 mM) > bromide (15-fold, *K*_d_ = 32 ± 2 mM) > nitrate (7.9-fold, *K*_d_ = 54 ± 4 mM) > iodide (3.8-fold, *K*_d_= 48 ± 7 mM) > sulfate (87%, *K*_d_ = 7.5 ± 1 mM) (Figures S7, S8, S16–S19, Table S5).

**Figure 5.**
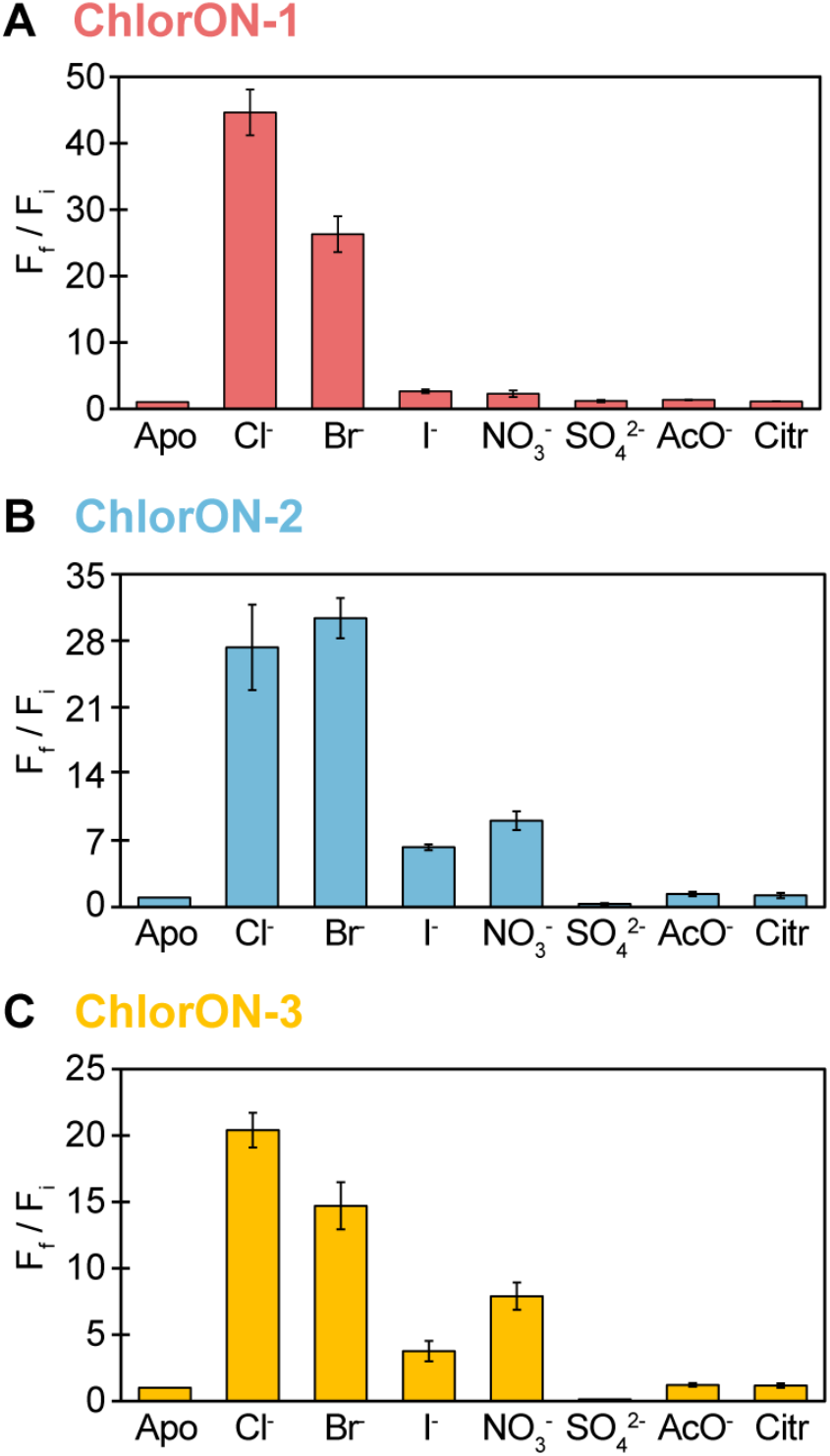
The anion selectivity profiles of the ChlorON sensors. The fluorescence response (F_f_/F_i_) of 10 μM (A) ChlorON-1, (B) ChlorON-2 and (C) ChlorON-3 to 1 mM (apo, F_i_) and 197 mM (F_f_) sodium chloride (Cl^-^), bromide (Br^-^), iodide (I^-^), nitrate (NO_3_^-^), sulfate (SO_4_^2-^), acetate (AcO^-^), and citrate (Citr) in 50 mM sodium phosphate buffer at pH 7 (λ_ex_ = 485 nm, λ_em_ = 515 nm). Titrations were carried out for anions with a response of 0.5 (turn-off) ≤ *F*_*f*_ */F*_*i*_ ≥ 2 (turn-on). The average of four technical replicates from two protein preparations with standard error of the mean is reported (Figure S13–S19, Table S5).

### Fluorescence imaging of labile chloride in FRT-CFTR cells with the ChlorONs

Encouraged by the robust *in vitro* properties of the ChlorONs, we next used live cell fluorescence imaging to determine if each sensor could provide a direct readout of chloride. It is important to note that of the anions tested *in vitro*, chloride is the most abundant, present at millimolar levels in extracellular media and cells.^1^ For these experiments, we selected the Fischer Rat Thyroid (FRT) cell model that endogenously expresses the sodium-iodide symporter (NIS) and stably overexpresses the cystic fibrosis transmembrane conductance regulator (CFTR).^89,90^ Activation of the CFTR channel with the diterpene forskolin (FSK) proceeds in a cyclic adenosine monophosphate (cAMP)-dependent manner, allowing for the transport of not only chloride but also iodide down the concentration gradient.^91^ Since the FRT cells do not express cAMP-dependent chloride channels, CFTR activity can be directly monitored using anion exchange assays.^89^

FRT-CFTR cells were transiently transfected with plasmids encoding non-targeted ChlorON sensors (Figure S20). After three days, cells were initially imaged in buffer containing 137 mM NaCl. The fluorescence signal was recorded from the same fields of cells during the time-lapse acquisition. To our surprise, cells expressing each sensor show a bright, intracellular fluorescence signal that is localized to the cytoplasm and nucleus (Figure 6 A–D, Figure S21). This led us to speculate if the ChlorON sensors could report on labile chloride pools. To test this, we relied on the fact that each sensor is more sensitive to chloride than iodide and carried out an exchange assay. The cells were perfused with buffer substituted with 100 mM NaI, 37 mM NaCl, and 20 μM FSK to trigger the efflux of chloride through CFTR and the influx of iodide through CFTR and NIS. As a result, the fluorescence signal is rapidly quenched by ∼59%, ∼43%, and ∼51% for ChlorON-1, ChlorON-2, and ChlorON-3, respectively (Figure 6E). Re-perfusion of the buffer containing 137 mM NaCl and 20 μM FSK increases the fluorescence signal by ∼2.1-fold for ChlorON-1, ∼1.5-fold for ChlorON-2, and ∼1.8-fold for ChlorON-3.

**Figure 6.**
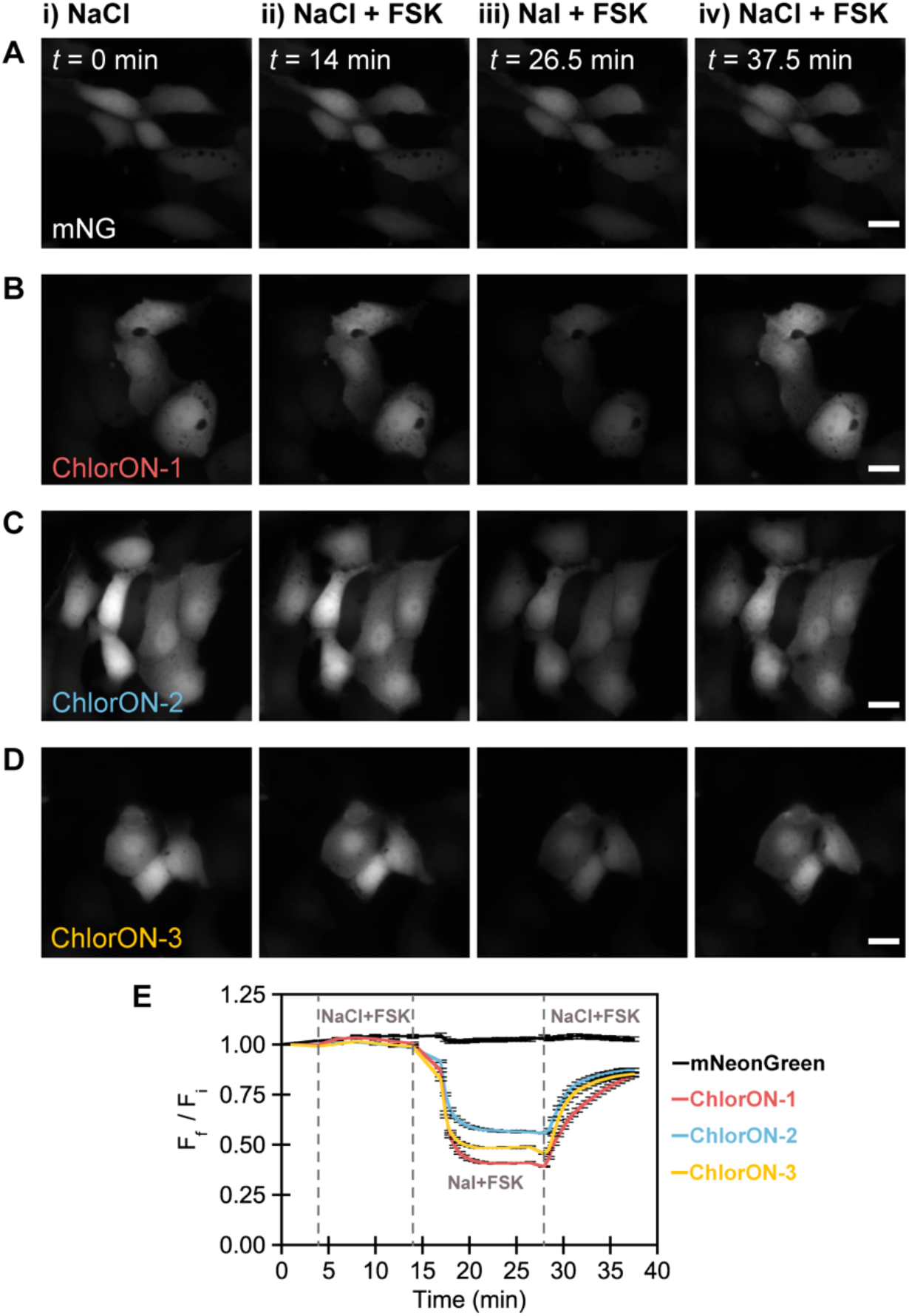
The ChlorON sensors provide a readout of labile chloride in live Fischer Rat Thyroid (FRT) cells overexpressing the cystic fibrosis transmembrane conductance regulator (CFTR). Representative fluorescence images of FRT-CFTR cells expressing (A) mNG, (B) ChlorON-1, (C) ChlorON-2, and (D) ChlorON-3 in imaging buffer supplemented with (i) 137 mM NaCl at *t* = 0 min (F_i_), (ii) 137 mM NaCl and 20 μM forskolin (FSK, F_f_) at *t* = 14 min, (iii) 100 mM NaI, 37 mM NaCl, and 20 μM FSK (F_f_) at *t* = 26.5 min, and (iv) 137 mM NaCl and 20 μM FSK (F_f_) at *t* = 37.5 min. Scale bar = 20 μm. Note: Exposure times are consistent within but variable between experiments. All experiments were carried out in an imaging buffer containing (2.7 mM KCl, 0.7 mM CaCl_2_, 1.1 mM MgCl_2_, 1.5 mM KH_2_PO_4_, 8.1 mM Na_2_HPO_4_, and 10 mM glucose) at pH 7.4 with the corresponding supplement listed in each panel. (E) Fluorescence response (F_f_/F_i_) of mNG (black line, n = 272 regions of interest (ROIs)), ChlorON-1 (red line, n = 256 ROIs), ChlorON-2 (blue line, n = 230 ROIs), and ChlorON-3 (yellow line, n = 255 ROIs). The vertical dashed lines correspond to the transition between each condition. The average fluorescence response (F_f_/F_i_) with standard error of the mean for all ROIs from three biological replicates is reported (Figure S21, Supplemental Video 1–4).

Finally, to confirm that the ChlorONs can directly report on changes in intracellular chloride, additional control experiments were performed under the same conditions with mNG and the pH-sensitive dye BCECF.^92,93^ Cells expressing the mNG parent have bright intracellular fluorescence, but no significant change (ΔF < 5%) is observed throughout the entire time-lapse acquisition (Figure 6). Moreover, as expected, the intracellular pH remains constant as indicated by the BCECF fluorescence (ΔF < 10%, Figure S22).^74,90,91^

## Discussion

In this study, we report the ChlorONs as the first single domain, turn-on fluorescent protein sensors for chloride that operate at physiological pH and in living cells. The ChlorONs were generated through a structure-guided approach by targeting the non-coordinating residues, K143 and R195, in mNG for combinatorial site-saturation mutagenesis. Since mNG is insensitive to chloride at physiological pH, a blind search was used to screen all possible amino acid combinations (19^2^ = 361) in *E. coli* lysate. To increase the probability of identifying chloride-sensitive variants *de novo*, bromide was first used as a surrogate anion. Here, we coin this approach as anion walking. The six variants resulting from our screen highlight how different combinations of residues at positions 143 and 195 could support a chloride binding pocket at physiological pH. At K143, aliphatic (glycine, leucine), aromatic (tryptophan, tyrosine), and charged (arginine) amino acids can be tolerated; at R195, mutations are enriched with isoleucine or leucine apart from one variant with lysine. The top three variants defining the ChlorON series – K143W/R195L (ChlorON-1), K143R/R195I (ChlorON-2), and K143R/R195L (ChlorON-3) – have overlapping substitutions that can be rationalized based on the *in vitro* spectroscopic properties.

At pH 7, all three sensors display a robust turn-on fluorescence response to chloride across a wide range of affinities. The observed sensing mechanism is linked to the stabilizing effect of chloride binding on the phenolate form of the chromophore (Figure 3). For ChlorON-1, this is more pronounced in the excited state. However, for ChlorON-2 and ChlorON-3, chloride tunes the phenolate chromophore between two different vibrational modes in the ground state, in addition to the excited state. Since R195L is common in both ChlorON-1 and ChlorON-3, the mutation at K143, as hypothesized, is key to tune the response through coordination. The side chains of both tryptophan and arginine could directly form electrostatic interactions with chloride or indirectly in a water-mediated fashion. Moreover, given that arginine is positively charged at physiological pH, the ion pairing interaction is effectively stronger. Thus, leading to the enhanced chloride binding affinities for ChlorON-2 and ChlorON-3 versus ChlorON-1.

Also, steric and hydrophobic effects from mutations at both positions cannot be ruled out. In combination with the described electronic effects, these factors could contribute to the size and shape of the binding pocket such that the desolvation and recognition of chloride is favored. Looking to the extreme end, K143W likely locks in the orientation of the residues in the coordination sphere. Chloride binding triggers a reorganization that translates to a greater perturbation of the excited state chromophore environment, which is reflected in the turn-on response for ChlorON-1 versus ChlorON-2 and ChlorON-3. These ideas can be extended to the anion selectivity profiles for each sensor which reveal a correlation between response and ionic radii. ChlorON-1 and ChlorON-3 both prefer chloride (1.81 Å) over bromide (1.96 Å) whereas ChlorON-2 has a comparable response to both. This difference can be linked to position 195. However, K143R unlocks the response of ChlorON-2 and ChlorON-3 to nitrate (1.79 Å) > iodide (2.20 Å) > sulfate (2.40 Å).^80,81^

To compensate for the mutations, new secondary interactions within and outside the binding pocket could also arise to affect the chromophore environment. This is clear in the pH-response profile of each sensor. Notably, ChlorON-2 and ChlorON-3 have two p*K*_a_s that increase with chloride. We attribute this to K143R since ChlorON-1 has one p*K*_a_ that can only be measured with chloride. The phenol form of the chromophore should be stabilized when the p*K*_a_ increases with chloride, suggesting a turn-off response like avYFP-H148Q, but this is not the case for all three ChlorONs.^54,55^ Furthermore, the magnitude of the response of each sensor is inversely correlated with the chloride binding affinity as the pH increases. It is possible that pH and chloride could tune the protonation state of nearby ionizable residues such as H62 and R88 that could form physical interactions with the chromophore.^88^ Such a stabilizing effect on the chromophore orientation would not necessarily be reflected in the p*K*_a_ but rather the fluorescence response to chloride. Looking forward, X-ray crystallography, ultrafast spectroscopy, and molecular dynamics simulations will be combined to fully capture these rich atomistic details.

Here, we have used fluorescence microscopy to show that the ChlorONs provide a reversible, direct readout of labile chloride in the FRT-CFTR cell model. By relying on the expression levels of the NIS and CFTR, all three sensors have a comparable response in the chloride-iodide exchange assay. This occurs despite differences in the measured binding affinities, suggesting that the comparable dynamic ranges across the low millimolar anion concentration regime could be a driving force for in-cell performance. While our present study highlights how the first-generation ChlorONs can be used to investigate chloride transport in living cells, we are excited by the untapped potential of this technology. Future efforts will integrate the power of protein engineering to improve sensor properties, including brightness, dynamic range, and sensitivity. We believe such an endeavor will transform the ChlorONs into a platform imaging technology, enabling researchers to monitor and measure chloride across time and space dimensions in new and unexplored biological contexts.

## Acknowledgements

The authors acknowledge members of the Dodani Lab for helpful discussions. The FRT-CFTR cells were kindly gifted by Dr. Jeong Hong from Emory University and the Cystic Fibrosis Foundation. S.C.D. acknowledges support for this work from UT Dallas, the Welch Foundation (AT-1918-20170325, AT-2060-20210327), and the National Institute of General Medical Sciences of the National Institutes of Health (R35GM128923). This study is the sole responsibility of the authors and does not represent the views of the funding sources.

## Author contributions

S.C.D. designed the research project. J.N.T. and W.S.Y.O. carried out the experiments. S.M.P., W.P., and H.G. provided technical assistance for pilot experiments. J.N.T., W.S.Y.O., and S.C.D. wrote the manuscript with input from all the authors.

## Safety statement

No hazardous chemicals or risks were encountered in this study. Standard biosafety level 1 and 2 procedures were followed for the appropriate experiments.

