## Supporting Information for "The ChlorON Series: Turn-on Fluorescent Protein Sensors for Imaging Labile Chloride in Living Cells"

### METHODS

**General.** All chemicals, reagents, and supplies were purchased from Integra Biosciences, Research Product International, Sigma-Aldrich, Thermo Fisher Scientific, VWR, or USA Scientific, unless otherwise noted. The protein structures and chemical structures in Figure 1 and Figure S1 were generated using MacPyMOL and ChemDraw (v 18.1), respectively.

**Bacterial plasmid design, construction, and preparation.** The gene encoding mNeonGreen (mNG, UniProt ID: A0A1S4NYF2) was codon optimized for expression in *Escherichia coli*, synthesized, and cloned into the pET-28b(+) vector between the NdeI and BamHI restriction sites with an N-terminal polyhistidine-tag as previously described (GenScript, Figure S2).[1] We note that the ten amino acids following the NdeI restriction site are not included in the numbering of the amino acid sequence (Figure S2).

The commercially prepared plasmid (4 ng) was reconstituted in 20 µL of autoclaved water, and the resulting 200 ng/µL stock solution was further diluted to 5 ng/µL with autoclaved water. One microliter of the diluted plasmid was used to transform *E. cloni* 10G ELITE Competent Cells (Lucigen) by electroporation (Bio-Rad Laboratories). Following this, the transformation mixture was plated on Miller's Luria Broth (LB, 10 g/L NaCl) agar plates containing 50 µg/mL kanamycin sulfate and incubated for ~18 h at 37 °C (New Brunswick Innova 42R, Eppendorf). A single colony was picked into 5 mL of LB containing 50 µg/mL kanamycin sulfate and incubated overnight for ~18 h at 37 °C with shaking at 250 rpm. The next day, the cells were harvested by centrifugation at 2,500*g* (5810 R, Eppendorf) for 5 min and stored at -20 °C. The plasmid was isolated using the QIAprep Spin Miniprep Kit (Qiagen) according to the manufacturer's instructions. The concentration of the purified plasmid was measured using the NanoDropLite Spectrophotometer (Thermo Fisher Scientific) and diluted to 10 ng/µL with autoclaved water for further use.

**Cloning of the double site-saturation mutagenesis library.** The double site-saturation mutagenesis library was generated using the mNG template prepared above. Two sets of four primers (3 forward and 1 reverse) were designed for the K143 and R195 mutation sites. Following the 22c-trick, each set of forward primers contained NDT, VHG, or TGG at the mutation site (Sigma-Aldrich, Table S1).[2] Each forward primer was dissolved in autoclaved water to a concentration of 100 µM, and the NDT, VHG, and TGG forward primers were combined in a ratio of 12:9:1 and diluted to a final concentration of 10 µM with autoclaved water. The reverse primers were diluted with autoclaved water to a final concentration of 10 µM. The polymerase chain reaction (PCR) was carried out using the Phusion High Fidelity PCR Kit (New England Biolabs) according to manufacturer's instructions. The reaction components and PCR conditions are shown in Table S2. The template DNA was removed by adding 1 µL DpnI (New England Biolabs) to the 50 µL PCR product and incubated for 2 h at 37 °C. Next, the PCR product was purified using agarose (Gold Biotechnology) gel electrophoresis and extracted using the Zymoclean Gel DNA Recovery Kit (Zymo Research). The concentration of the purified PCR product was determined using the NanoDropLite Spectrophotometer and diluted to 10 ng/µL with CutSmart Buffer (New England Biolabs) to a volume of 10 µL. This diluted PCR product was combined with 10 µL of the Gibson Assembly Master Mix (New England Biolabs) and purified using the DNA Clean & Concentrator Kit (Zymo Research) according to manufacturer's instructions. *E. cloni* EXPRESS BL21(DE3) Competent Cells (Lucigen) were transformed with one microliter of the purified DNA via electroporation. The resulting transformation mixture was plated on LB agar plates containing 50 µg/mL kanamycin sulfate as described above. To verify that both sites were successfully mutated, the plasmids from five random colonies were prepared and isolated as described above for Sanger sequencing (Eurofins Scientific).

**Library expression, screening, and validation.** *E. cloni* EXPRESS BL21(DE3) Competent Cells were transformed with the plasmid encoding the mNG parent as described above. For the mNG parent plate, eight colonies of mNG were picked into 500 µL of modified 2xYT media (16 g/L Bacto Tryptone,10 g/L yeast extract, and 50 µg/mL kanamycin sulfate) in a 96-well deep well plate (Greiner Bio-One). For the library, 94 colonies were picked into a separate 96-well deep well plate with each well containing 500 µL of the modified 2xYT media. For each 96-well plate, two wells contained media only to check for contamination. This process was repeated until twenty 96-well deep well plates were picked with a total of ~1,880 colonies from the library. Each plate was sealed with an Easy App microporous film (USA Scientific) and incubated for ~18 h at 37 °C with shaking at 200 rpm (New Brunswick Innova 44R, Eppendorf). The next day, each well of a 96-well deep well plate was filled with 950 µL of the modified 2xYT media and inoculated with 50 µL of the overnight culture using a Biomek NXP liquid handler (Beckman Coulter). After sealing, the plates were incubated for 2.5 h at 37 °C with shaking at 200 rpm. Protein expression was induced with the addition of 50 µL of 21 mM isopropyl β-D-thiogalactopyranoside (IPTG, Gold Biotechnology) in the modified 2xYT media to a final concentration of 1 mM. The plates were further incubated for ~22 h at 37 °C with shaking at 200 rpm and then incubated at 10 °C for at least 8 h without shaking. The cells were harvested by centrifugation at 2,500*g* (5810 R, Eppendorf) for 15 min and stored as pellets at -20 °C.

On the day of the screen, the plates were subjected to three freeze-thaw cycles for 15 min at room temperature. Then, each well was resuspended in 500 µL of cold lysis buffer (25 mM sodium phosphate buffer at pH 8 containing 2 mg/mL lysozyme, 10 µg/mL deoxyribonuclease I (DNase I), and 2 mM $MgCl_2$) and gently vortexed until all cell pellets were dislodged (Vortex 2 Shaker, IKA Works). The plates were further incubated for 1 h at 37 °C with shaking at 200 rpm. Following this, the cell lysates were clarified by centrifugation at 2,500*g* for 30 min at 4 °C, and 175 µL of the supernatant was transferred to a clear 96-well microtiter plate (Caplugs) using the liquid handler. For each well, excitation was provided at 485 nm (5 nm bandwidth), and the emission intensity was measured at 520 nm ($F_i$, 5 nm bandwidth, 30 flashes, 60 gain) on a Spark 10M plate reader (Tecan) at room temperature (22–25 °C). Following these baseline measurements, 25 µL of 25 mM phosphate buffer at pH 8 containing 0.8 M NaBr was added to each well to a final concentration of 100 mM NaBr, and the emission intensity at 520 nm was recorded again ($F_f$, Figure S3). The mNG parent plate was tested first to determine the average fluorescence response with standard error of the mean ($F_f/F_i$ = 1.2 ± 0.01) and coefficient of variance (~2%) before proceeding with the library plates. Variants from the library parents with at least a 4-fold turn-on fluorescence response ($F_f/F_i$) to bromide were freshly restreaked from the overnight cultures onto LB agar plates containing 50 µg/mL kanamycin sulfate and incubated overnight at 37 °C for rescreening.

For each restreaked variant, two colonies were picked into 5 mL of the modified 2xYT media and incubated for ~18 h at 37 °C with shaking at 250 rpm. The next day, 25 mL of the modified 2xYT media in 125-mL baffled flasks were inoculated with 500 µL of the overnight culture and incubated at 37 °C with shaking at 250 rpm. After 2.5 h, protein expression was induced by adding 25 µL of 1 M IPTG in water to a final concentration of 1 mM. The cell cultures were incubated for ~22 h at 37 °C with shaking at 250 rpm, then harvested via centrifugation at 2,500*g* for 5 min at 4 °C (5810 R, Eppendorf), and finally stored as pellets at -20 °C. The following day, each cell pellet was thawed on ice for 30 min, resuspended in 2 mL of 15 mM sodium phosphate buffer at pH 7.5, and transferred to a 2-mL centrifuge tube. The cells were then lysed via sonication on ice at 30% amplitude with a 20 s/40 s on-off pulse sequence for 2 min (Q500, QSonica) and clarified by centrifugation at 20,000*g* for 20 min at 4 °C (5424 R, Eppendorf). For each biological replicate, 250 µL of the supernatant was diluted into 750 µL of 25 mM sodium phosphate buffer at pH 7 or pH 8. A 25 µL portion of the diluted cell lysate was added to 175 µL of 25 mM sodium phosphate buffer containing 0 mM or 114 mM NaCl at pH 7 or 8 to a final concentration of 0 mM ($F_i$) or 100 mM ($F_f$) NaCl in a clear 96-well microtiter plate. For each well, excitation was provided at 485 nm (5 nm bandwidth), and the emission was collected from 505–650 nm (5 nm step size, 5 nm bandwidth, 30 flashes, 100 gain). The emission intensity at 520 nm was used to calculate the turn-on fluorescence response ($F_f/F_i$). The average of the two biological replicates with standard error of the mean is reported (Table S3). In parallel, the plasmids were isolated from each biological replicate using the QIAprep Spin Miniprep Kit for Sanger sequencing (Eurofins Scientific, Table S3). These data were used to identify the top three variants with turn-on fluorescence responses $F_f/F_i \leq$ 25-fold with 100 mM NaCl at pH 7 for further characterization in purified form: K143W/R195L (ChlorON-1), K143R/R195I (ChlorON-2), and K143R/R195L (ChlorON-3).

**Large-scale protein expression and purification.** The sequence-verified plasmids encoding mNG, ChlorON-1, ChlorON-2, and ChlorON-3 were transformed into *E. cloni* EXPRESS BL21(DE3) Competent Cells and selected on LB agar plates as described above. Single colonies were picked into 25 mL of 2xYT media (16 g/L Bacto Tryptone, 10 g/L yeast extract, 5 g/L NaCl, and 50 µg/mL kanamycin sulfate) in 125-mL baffled flasks and incubated overnight at 37 °C with shaking at 250 rpm. The next day, 600 mL of 2xYT media in 2-L baffled flasks were inoculated with 24 mL of the overnight culture and incubated for 2 h at 37 °C with shaking at 150 rpm until the $OD_{600}$ reached ~0.6–0.8. Protein expression was induced with 600 µL of 1 M IPTG in water to a final concentration of 1 mM. Then, the cultures were incubated with shaking at 150 rpm as follows: 20–22 h at 37 °C for mNG and ChlorON-1, 48 h at 18 °C for ChlorON-2, and 24 h at 23 °C for ChlorON-3. Following this, the cultures were incubated at 10 ºC for 2 h, then harvested by centrifugation at 3,200*g* for 35 min at 4 °C (5810 R, Eppendorf), and resuspended in ~30 mL of 20 mM Tris buffer at pH 7.5 containing 200 mM NaCl, 5 mM $MgCl_2$, 30 µg/mL DNase I, and protease inhibitor (1 protease inhibitor capsule/500 mL, Pierce). The resuspended cells were stored at -20 °C until purification.

For protein purification, the cell suspension was thawed overnight at 10 °C and lysed the following day by sonication on ice at 30% amplitude with a 15 s/45 s on-off pulse sequence for 5 min. The cell lysate was clarified by ultracentrifugation at 18,000*g* for 30 min at 4 °C (Optima XPN-80, Beckman Coulter). The following purification steps were carried out at 4 °C. Prior to sample loading, a 5-mL nickel nitrilotriacetic acid column (Nuvia IMAC, Bio-Rad Laboratories) was equilibrated with 5 column volumes (CV) of running buffer (20 mM Tris buffer at pH 7.5 containing 200 mM NaCl and 30 mM imidazole) using the NGC Quest 10 Chromatography System (Bio-Rad Laboratories). Following this, the clarified supernatant was loaded using a sample pump at a flow rate of 4 mL/min. The column was first washed with 10 CV of the running buffer at a flow rate of 4 mL/min, and the His-tagged protein was eluted with a 0–100% gradient of the running buffer and elution buffer (20 mM Tris buffer at pH 7.5 containing 200 mM NaCl and 500 mM imidazole) at a flow rate of 5 mL/min for 20 CV. The absorbance intensities at 280 nm and 480 nm were monitored, and the fractions within the chromatogram peak that absorbed at both wavelengths were pooled and loaded onto a pre-equilibrated desalting column (HiPrep 26/10 Desalting, Cytiva). The protein sample was eluted using an exchange buffer (20 mM Tris buffer at pH 7.5 containing

200 mM NaCl) at a flow rate of 4 mL/min. The eluted fractions that absorbed at 280 nm and 480 nm were pooled and concentrated to a final volume of ~13 mL using a 15-mL Amicon filter with a molecular weight cut-off (MWCO) of 10 kilodalton (kDa, MilliporeSigma). Then, the protein was loaded onto a pre-equilibrated size exclusion chromatography column (Hi-Load Superdex 26/600 200 prep grade, Cytiva) and eluted using the exchange buffer at a flow rate of 2.6 mL/min. The fractions with absorbance intensities at both 280 nm and 480 nm that eluted at ~0.7 CV corresponding to the monomeric protein (~26.6 kDa) based on a known protein standard (Bio-Rad Laboratories) were collected and buffer exchanged into 20 mM sodium phosphate buffer at pH 7.4 containing 50 mM NaCl using centrifugal filters with a 10-kDa MWCO (Figure S4A). For each protein, two biological replicates were independently expressed, purified, and characterized as further described below.

**SDS-PAGE.** To evaluate the purity of each protein preparation, stock solutions were diluted to ~0.3 mg/mL and combined with 4X Laemmli Buffer (Bio-Rad Laboratories) containing 10% β-mercaptoethanol in a 3:1 (v/v) ratio. Each sample was boiled for 5 min at 95 °C with shaking at 300 rpm (ThermoBlock, Eppendorf), and then 15 µL of each sample was loaded onto a 12% TGX FastCast acrylamide gel alongside 4 µL of the Precision Plus Protein unstained standard (Bio-Rad Laboratories). The electrophoresis was carried out using a Mini-PROTEAN Tetra Cell setup and a PowerPac power supply (Bio-Rad Laboratories) at 240 mV for ~30 min in 1X Tris-Glycine-SDS buffer. The gel was visualized as previously described using a Coomassie stain (Figure S4B).[3]

**Protein concentration determination.** To determine the protein concentration, each aliquot of purified protein was diluted 50-fold into 50 mM sodium phosphate buffer at pH 7. This solution was then diluted 1.25-fold into the same buffer and serially diluted seven more times to generate a standard curve. A portion of each solution (200 µL) was transferred to a 96-well UV-Star microtiter plate (Greiner Bio-One). Absorbance spectra were collected from 250–350 nm with a 3.5 nm bandwidth using the plate reader. The corrected absorbance intensities were calculated using Equation 1:

$$A_{\text{corrected}} = A_{280} - A_{320} \tag{1}$$

where ($A_{280}$) corresponds to the absorbance intensity at 280 nm for the protein and ($A_{320}$) corresponds to absorbance intensity at 320 nm as a baseline. The $A_{corrected}$ was plotted versus the dilution factor of the protein samples to obtain a slope ($G_s$). The absorbance intensities that fell outside the linear portion of the curve (>0.2) were excluded from the analysis. From this, the slope ($G_s$) was used in Equation 2 to determine the absorbance ($A$) of the stock solution:

$$A = G_s * d \tag{2}$$

where $d$ corresponds to the initial dilution factor of 50. Using this calculated absorbance value ($A$), the Beer-Lambert law was used to determine the protein concentration ($c$) with Equation 3:

$$c = \frac{A}{\varepsilon * l} \tag{3}$$

where $l$ corresponds to the optical pathlength and $\varepsilon$ corresponds to the extinction coefficient at 280 nm. The $\varepsilon$ values for ChlorON-1 (49,850 $M^{-1}cm^{-1}$), ChlorON-2 (44,350 $M^{-1}cm^{-1}$), and ChlorON-3 (44,350 $M^{-}$

$^{1}$cm$^{-1}$) were determined using the ProtParam tool in ExPasy.[4] The optical pathlength ($l$) was determined by collecting the absorbance intensities at 975 nm ($A_{975}$) and 900 nm ($A_{900}$) of 200 µL water in a 96-well UV-Star microtiter plate on a plate reader (3.5 nm bandwidth) and 3 mL water in a quartz cuvette (1 cm pathlength, Hellma USA) on a UV-vis spectrophotometer (2 nm bandwidth, Agilent).[5] The pathlength for the microtiter plate ($l$) was determined to be ~0.6 cm using Equation 4:

$$l = \frac{A_{975(\text{well})} - A_{900(\text{well})}}{K\text{-}factor} * 1\ \text{cm} \tag{4}$$

where the *K-factor* is a constant that corresponds to the difference between $A_{975}$ and $A_{900}$ in the cuvette (*K-factor* = 1.68). Based on this, the purified proteins were concentrated to ~500 µM and stored at -20 °C until further use.

**General spectroscopy methods.** Protein aliquots were thawed on ice prior to testing. For all measurements except the extinction coefficients and quantum yields, the following plate reader settings were used. All measurements were carried out at room temperature (22–25 °C). Absorbance spectra were collected from 300–650 nm (5 nm step size, 3.5 nm bandwidth). For the excitation provided at 400 nm (5 nm bandwidth), the emission was collected from 425–650 nm (5 nm step size, 5 nm bandwidth, 30 flashes, 150 gain). For the excitation provided at 485 nm (5 nm bandwidth), the emission was collected from 500–650 nm (5 nm step size, 5 nm bandwidth, 30 flashes, 110 gain). For each protein preparation ($n$ = 2), two technical replicates were carried out for all measurements.

**Chloride titrations and apparent dissociation constants.** Aliquots of purified protein were diluted 50-fold to ~10 µM in 50 mM sodium phosphate buffer at pH 6, 7, or 8 containing 0, 1, 3, 6, 12, 25, 50, 100, 200, or 400 mM NaCl to a final concentration of 1, 2, 3.9, 6.9, 13.3, 25.5, 50, 99, 197, or 393 mM, respectively. A portion of each sample (100 µL) was transferred to two wells of a 96-well half-area microtiter plate (Greiner Bio-One) for plate reader measurements.

The average fluorescence intensity and standard error of the mean ($\sigma_M$) were used to calculate the average emission response ($F_f/F_i$) where $F_i$ corresponds to the fluorescence intensity with 1 mM chloride and $F_f$ corresponds to the fluorescence intensity with each chloride concentration tested. These values were used to calculate the propagated error of the mean ($\sigma$) for the average emission responses of each protein preparation using Equation 5:

$$\sigma = \frac{F_f}{F_i} * \sqrt{\left(\frac{\sigma_{M-F_f}}{F_f}\right)^2 + \left(\frac{\sigma_{M-F_i}}{F_i}\right)^2} \tag{5}$$

The propagated errors of the mean ($\sigma$) from Equation 5 for both protein preparations were used to determine the grouped data standard deviation ($SD$) using Equation 6:

$$SD = \sqrt{\frac{1}{N-1}\left(\sum_{i=1}^{j}(n-1)_i\,\sigma_i^2 + \sum_{i=1}^{j} n_i(\bar{x}_i - \bar{x})^2\right)} \tag{6}$$

where $i$ corresponds to the index of summation for each protein preparation, $N$ corresponds to the number of technical replicates from both protein preparations ($N$ = 4), $n$ corresponds to the number of replicates for each protein preparation ($n$ = 2), $\bar{x}_i$ corresponds to the average emission response ($F_f/F_i$) of the all technical replicates for each protein preparation ($n$ = 2), and $\bar{x}$ corresponds to the average emission response ($F_f/F_i$) for all replicates ($N$ = 4) (Figure 3 and Table S4).

For each protein preparation, the apparent dissociation constant ($K_d$) was determined by plotting the average fluorescence intensity at 515 nm ($\lambda_{ex}$ = 485 nm) with standard error of the mean ($\sigma_M$) versus the chloride concentration [$Cl^-$] in KaleidaGraph v4.5 (Synergy Software) using Equation 7:

$$F_{\mathrm{obs}} = \frac{[Cl^-] * (F_{\max} - F_{\min})}{(K_d + [Cl^-])} + F_{\min} \tag{7}$$

where $F_{obs}$ is the average fluorescence intensity at each concentration tested, and $F_{min}$ and $F_{max}$ are the average fluorescence intensities in the presence of 1 mM and 393 mM NaCl, respectively.

The grouped data standard deviation ($SD$) for both protein preparations was determined using Equation 6 where $\sigma$ corresponds to the error of each fit from KaleidaGraph, $\bar{x}_i$ corresponds to the average $K_d$ of the two technical replicates for each protein preparation, and $\bar{x}$ corresponds to the average $K_d$ for all four replicates (Figure 3, Figure S6–S8, and Table S4).

**Protein extinction coefficients and quantum yields.** To determine the extinction coefficients ($\varepsilon$), aliquots of purified protein were diluted 50-fold to ~10 µM in 50 mM sodium phosphate buffer at pH 7. These solutions were diluted 1.25-fold into the same buffer and serially diluted seven more times to generate a standard curve. A portion of each solution (200 µL) was transferred to a 96-well UV-Star microtiter plate, and absorbance spectra were collected from 250–650 nm (2 nm step size, 3.5 nm bandwidth).

For each well, the absorbance intensities at 280 nm ($A_{280}$) for total protein and 480 nm ($A_{480}$) for the chromophorylated protein were plotted versus the sample dilution factors to obtain the slopes $\Delta A_{280}$ and $\Delta A_{480}$, respectively. Since the optical pathlengths ($l$) and protein concentrations ($c$) are constant at both wavelengths, the Beer-Lambert law in Equation 3 can be simplified to Equation 8:

$$\varepsilon_{480} = \varepsilon_{280} * \left(\frac{\Delta A_{480}}{\Delta A_{280}}\right) \tag{8}$$

Where $\varepsilon_{280}$ and $\varepsilon_{480}$ correspond to the extinction coefficients at 280 nm and 480 nm, respectively. The experiment was repeated for each protein in the presence of 197 mM NaCl. For both ChlorON-2 and ChlorON-3, there is an additional absorbance peak at 506 nm in the presence of 197 mM NaCl. The $\varepsilon$ at 506 nm ($\varepsilon_{506}$) was also determined. The average extinction coefficients of the four technical replicates from two protein preparations with the grouped data standard deviation (Equation 6) are reported (Table S4).

In parallel, the quantum yields for ChlorON-1, ChlorON-2, and ChlorON-3 ($\Phi_{FP}$) were determined with reference to fluorescein in 100 mM sodium hydroxide.[6] Briefly, excitation was provided at 488 nm (5 nm bandwidth) and 460 nm (5 nm bandwidth), and the emission was collected from 506–750 nm and 490–750 nm, respectively (2 nm step size, 5 nm bandwidth, 30 flashes, 75 gain). For each spectrum, the area under the curve was integrated using Excel. The ratio of the integrated areas from 506–750 nm for $\lambda_{ex}$ = 460 nm and $\lambda_{ex}$ = 488 nm was used to estimate the integrated area from 490–506 nm for the emission spectra of $\lambda_{ex}$ = 488 nm. Linear plots were generated by plotting the integrated area from 490–750 nm ($\lambda_{ex}$ = 488 nm) versus the absorbance intensities at 488 nm. Data points that fell outside the linear portion of the curve (absorbance intensities > 0.06) were excluded analysis. Based on this the quantum yields were determined using Equation 9:

$$\Phi_{\mathrm{FP}} = \Phi_{\mathrm{Ref}} * \left(\frac{Slope_{\mathrm{FP}}}{Slope_{\mathrm{Ref}}}\right) * \left(\frac{\eta_{\mathrm{FP}}}{\eta_{\mathrm{Ref}}}\right)^2 \quad (9)$$

where $\Phi_{\mathrm{Ref}}$ corresponds to quantum yield of fluorescein ($\Phi_{\mathrm{Ref}}$ = 0.92), $slope_{\mathrm{FP}}$ and $slope_{\mathrm{Ref}}$ correspond to the slope from the linear plots for protein and fluorescein, respectively, and $\eta_{FP}$ and $\eta_{Ref}$ correspond to the refractive index of water ($\eta$ = 1.33).[6,7] The average of four technical replicates from two protein preparations with the grouped data standard deviation (Equation 6) is reported (Figure S9 and Table S4).

**Chromophore p*K*<sub></sub>**

**Chromophore p$K_a$s.** Aliquots of purified protein were diluted 50-fold to ~10 µM protein in 50 mM sodium acetate buffer from pH 3.5–5.5 or in 50 mM sodium phosphate buffer from pH 5.5–8.5 containing 0 mM or 200 mM NaCl (1 mM or 197 mM final concentration, respectively). A portion of each sample (100 µL) was transferred to two wells of a 96-well half-area microtiter plate for plate reader measurements as described above (see *General spectroscopy methods*). For each protein preparation, the p$K_a$s were determined by plotting the average fluorescence intensities at 515 nm ($\lambda_{ex}$ = 485 nm) with standard errors of the mean ($\sigma_M$) versus the pH in Kaleidagraph. The p$K_a$s for ChlorON-1 were determined in the presence of 1 mM and 197 mM NaCl using the following equation:

$$F_{515} = \frac{a+\left(b*10^{(pH-pKa)}\right)}{1+10^{(pH-pKa)}} \quad (10)$$

where *a* corresponds to the average minimum fluorescence intensity at acidic pH and *b* corresponds to average maximum fluorescence intensity at basic pH.[8] Since the emission response curves for ChlorON-2 and ChlorON-3 are bell-shaped, Equation 10 was modified to Equation 11 to fit the data to two different p$K_a$s as follows:

$$F_{515} = \frac{a_1+\left(b_1*10^{(pH-pKa_1)}\right)}{1+10^{(pH-pKa_1)}} * \frac{a_2+\left(b_2*10^{(pH-pKa_2)}\right)}{1+10^{(pH-pKa_2)}} \quad (11)$$

where $a_1$ and $b_1$ correspond to the average minimum fluorescence intensity at acidic pH and the average maximum fluorescence intensity at a higher or neutral pH, respectively, and $a_2$ and $b_2$ correspond to the average maximum fluorescence intensity at acidic or neutral pH and the average minimum fluorescence intensity at basic pH, respectively.[8] The average of four technical replicates from two protein preparations with the grouped data standard deviation (Equation 6) is reported (Figure 4, Figure S10–S12, and Table S4).

**Anion selectivity.** Absorbance and fluorescence spectra were collected as described above in the *General spectroscopy methods*. Aliquots of purified mNG, ChlorON-1, ChlorON-2, and ChlorON-3 were diluted 50-fold to ~10 µM in 50 mM phosphate buffer at pH 7 containing 0 mM or 200 mM sodium chloride, bromide, iodide, nitrate, sulfate, acetate, or citrate (0 mM or 196 mM final concentration, respectively) (Figure 5). For phosphate, aliquots of the purified proteins were diluted 50-fold to ~10 µM in 50 mM HEPES buffer at pH 7 containing 0 mM or 200 mM sodium phosphate (0.4 mM or 196 mM final concentration, respectively) (Figure S13). Given the original protein stocks, all solutions contained a final concentration of 1 mM sodium chloride. The average fluorescence intensity and standard error of the mean ($\sigma_M$) were used to calculate the average emission response ($F_f/F_i$) where $F_i$ corresponds to the fluorescence intensity with 0 mM anion and $F_f$ corresponds to the fluorescence intensity with 196 mM anion (Figure S14–S19). For a given anion, if 0.5 (turn-off) $\leq F_f/F_i \geq$ 2 (turn-on), the $K_d$ was determined. To do this, anion titrations were carried out in 50 mM sodium phosphate buffer at pH 7 containing 0, 1, 3, 6, 12, 25, 50, 100, 200, or 400 mM sodium bromide, iodide, nitrate, or sulfate (0, 1.0, 2.9, 5.9, 12.3, 24.5, 49, 98, 196, and 392 mM final concentration, respectively). The $K_d$ for each sensor was determined as described above (Equation 7). The average of four technical replicates from two protein preparations with the grouped data standard deviation (Equation 6) is reported (Figure 4, Figure S13–S19, and Table S5).

**FRT-CFTR cell culture.** Fischer rat thyroid cells (FRT) stably expressing the cystic fibrosis transmembrane conductance regulator (CFTR) were provided by Dr. Jeong Hong from Emory University and the Cystic Fibrosis Foundation and cultured as previously described.[9] Briefly, cells were grown in Ham's F-12 with Coon's modification media (Sigma-Aldrich) containing 10% fetal bovine serum (FBS), 100 µg/mL hygromycin B, and 100 µg/mL zeocin at 37 °C, 5% $CO_2$. Cells were maintained in a T25 flask (Corning) and were split at regular intervals by washing twice with 5 mL of 1X Phosphate Buffered Saline (PBS, Gibco), followed by the addition of 3 mL trypsin-EDTA (0.05%) (Gibco) for 20 min at 37 °C, 5% $CO_2$. The trypsin reaction was quenched with 6 mL of the Ham's F-12 media described above. Following this, the cells were harvested via centrifugation at 200*g* (5702, Eppendorf) for 5 min and resuspended in 3 mL of fresh media. For cell counting, 2 µL of the resuspended cells were diluted into 10 µL of 0.4% trypan blue (Sigma-Aldrich) in PBS and 8 µL of media in a 500 µL centrifuge tube. The resulting suspension was pipetted into a hemocytometer for analysis.

**Fluorescence imaging and analysis of FRT-CFTR cells transfected with mNG and ChlorON-1/2/3.** Cell transfections were conducted using the genes encoding for mNG, ChlorON-1, ChlorON-2, and ChlorON-3. All genes were codon optimized for expression in mammalian cells and cloned into the pcDNA3.1(+)-N-6His vector between the BamHI and EcoRI restriction sites (GenScript, Figure S20). Technical grade plasmid preparations (100 µg/mL) were purchased and used for transfections (GenScript). For all transfections, 1 µg of each plasmid was complexed with 1.5 µL of the Lipofectamine 3000 reagent and 2 µL of the P3000 enhancer reagent (Invitrogen) in 250 µL of Reduced Serum Opti-MEM (Gibco) according to manufacturer's instructions. After 20 min, the complex was seeded in a 35-mm dish with 10-mm glass coverslip (No. 1.5, Cellvis), followed by plating of 2.5 x $10^5$ cells. Each dish was incubated for three days at 37 °C, 5% $CO_2$ prior to imaging with fluorescence microscopy.

Aliquots of the CFTR agonist forskolin (FSK, 20 mM, Sigma-Aldrich) were prepared in dimethylsulfoxide (DMSO) and frozen until further use. On the day of the experiment, the media was aspirated, and the cells were washed with 2 mL of a modified PBS buffer (2.7 mM KCl, 0.7 mM $CaCl_2$, 1.1 mM $MgCl_2$, 1.5 mM $KH_2PO_4$, 8.1 mM $Na_2HPO_4$, and 10 mM glucose at pH 7.4) supplemented with 137 mM NaCl, followed by incubation for 30 min at 37 °C in a stage top incubator equipped with an automated perfusion system (Tokai Hit). The stage top incubator and perfusion system were kept at 37 °C throughout the imaging experiment. Differential interference contrast (DIC) and fluorescence images were collected using a 20X air objective with a numerical aperture of 0.7 on an inverted fluorescence microscope (IX83, Olympus) and a FLED LED System set to level 1 (Sutter). The EGFP/FITC/Cy2 excitation filter centered at 470 nm (40 nm bandwidth, Chroma) was used with an EGFP emission filter centered at 525 nm (50 nm bandwidth, Chroma). Exposure was provided at 25% from 20–60 ms for mNG and at 100% from 150–600 ms for ChlorON-1/2/3. For each biological replicate, two different fields were selected, and the x, y, and z coordinates were recorded to image the same fields after each treatment using the CellSens software. The Z-drift compensation (ZDC) was set in Single Shot Mode to maintain the focal plane during acquisition. At the start of each experiment, time-lapse images were collected every 2 min for 4 min to establish the baseline fluorescence signal. Then, the imaging solution was then exchanged with the modified PBS buffer supplemented with 137 mM NaCl and 20 µM FSK at a rate of 4 mL/min for 2 min. Following this, images were captured every 2 min for 10 min. In the next treatment, the modified PBS buffer supplemented with 100 mM NaI, 37 mM NaCl, and 20 µM FSK was perfused at 4 mL/min for 2 min, and after a 1 min delay, time-lapse images were collected every 30 s for 10 min. The modified PBS buffer supplemented with 137 mM NaCl and 20 µM FSK was then re-perfused at 4 mL/min for 2 min, and after a 1 min delay, images were captured every 30 s for 10 min.

For the analysis, the Fiji is Just ImageJ (Fiji v2.0) software was used. All time-lapse images were first concatenated into a single stack.[10] The fluorescence channel was then sharpened, and the StackReg plugin with the transformation set to Translation was used to align each slice in the stack.[11] The auto default threshold was used on the maximum intensity Z-projection to create a mask. From this mask, the Fiji Analyze Particles function was used to select regions of interest (ROIs) corresponding to cells greater than 20 pixels in length with a circularity of 0–1. Cells in close contact that could not be differentiated by the software were considered as one ROI. ROIs with saturated intensity or debris were manually excluded from the selection. The ROIs were then transferred to the raw fluorescence images for each field, and the median fluorescence intensity was measured for each time point using the Multi Measure function in Fiji. The fluorescence intensity for each ROI ($F_f$) was normalized to the initial fluorescence intensity ($F_i$) of the same ROI. At least two different fields were sampled for each biological replicate, and the average turn-on response ($F_f/F_i$) with standard error of the mean is reported for all ROIs from three biological replicates (Figure 6, Figure S21, and Supplemental Video 1–4).

**BCECF staining, imaging, and analysis.** Aliquots of the intracellular pH indicator 2',7'-bis-(2-carboxyethyl)-5-(6)-carboxyfluorescein-acetoxymethyl ester (BCECF-AM, 1 mM, Invitrogen) were prepared in anhydrous DMSO and frozen until further use. Each 35-mm imaging dish was seeded with 2.5 x $10^5$ cells and incubated for three days at 37 °C, 5% $CO_2$ as described above. On the day of the experiment, the media was aspirated, and the cells were washed once with 2 mL of Dulbecco's Modified Eagle's Medium (DMEM) formulated with 4.5 g/L glucose and 110 mg/L sodium pyruvate. The cells were then stained with 2 mL of DMEM containing 5 µM BCECF-AM and incubated for 1 h at 37 °C, 5% $CO_2$. The BCECF-AM dye is nonfluorescent until it is hydrolyzed by intracellular esterases to the fluorescent BCECF acid form.[12,13] After staining, the cells were washed twice with 2 mL of 1X PBS at pH 7.4 (Gibco) and then washed twice with 2 mL of a modified PBS buffer (2.7 mM KCl, 0.7 mM $CaCl_2$, 1.1 mM $MgCl_2$, 1.5 mM $KH_2PO_4$, 8.1 mM $Na_2HPO_4$, and 10 mM glucose at pH 7.4) supplemented with 137 mM NaCl. Following this, the cells were incubated for 30 min at 37 °C in the stage top incubator prior to imaging.

Throughout the imaging experiment, the stage top incubator and perfusion system were kept at 37 °C. BCECF was excited with two excitation filters centered at 495 nm (10 nm bandwidth, Chroma) and at 436 nm (20 nm bandwidth, Chroma) that correspond to the pH-sensitive and pH-insensitive absorption maxima, respectively.[12,13] For both excitations, an emission filter centered at 540 nm (40 nm bandwidth, Chroma) was used. The exposure was provided at 25% for 200 ms. For each dish, the position coordinates of two different fields were selected in the CellSens software as described above, and the ZDC was set in Single Shot Mode to maintain focus. At the beginning of the experiment, baseline fluorescence images were collected every 2 min for 4 min. Then, the modified PBS buffer supplemented with 137 mM NaCl and 20 µM FSK was perfused at 4 mL/min for 2 min as described above, and images were captured every 2 min for 10 min. Next, the modified PBS buffer supplemented with 100 mM NaI, 37 mM NaCl, and 20 µM FSK was perfused, and after a 1 min delay, images were collected every 1 min for 10 min. Following this, the imaging solution was exchanged with the modified PBS buffer supplemented with 137 mM NaCl and 20 µM FSK, and after a 1 min delay, the cells were imaged every 1 min for 15 min. At the end of the experiment, the imaging solution was manually replaced on the stage with a pH clamping buffer at pH 8 (137 mM KCl, 2.7 mM NaCl, 0.7 mM $CaCl_2$, 1.1 mM $MgCl_2$, 1.5 mM $KH_2PO_4$, and 8.1 mM $Na_2HPO_4$) supplemented with 5 µM valinomycin (Sigma-Aldrich) and 5 µM nigericin (Sigma-Aldrich). After a 5 min incubation period, the same fields were imaged every 2 min for 4 min.

The imaging analysis was carried out using the Fiji software described above. For each BCECF fluorescence channel, all time-lapse images were concatenated into a single stack, sharpened, and aligned using the StackReg plugin in Translation mode.[10,11] For the BCECF dye, an additional step of background subtraction (50 pixels) was applied to each fluorescence channel. The background subtracted BCECF fluorescence ($\lambda_{ex}$ = 495 nm) images were used to find the maximum intensity Z-projection, and these images were then auto thresholded and used to create a mask and select the ROIs. Cells in close contact that could not be differentiated by the software were considered as one ROI. ROIs with saturated intensity or debris were manually excluded from the selection. For each field, the ROIs were then transferred to the background subtracted fluorescence images with $\lambda_{ex}$ = 495 nm ($F_{Ex495}$) and $\lambda_{ex}$ = 436 nm ($F_{Ex436}$), and the median fluorescence intensity was measured for each time point in Fiji. The BECEF fluorescence intensity ratio ($F_{Ex495}/F_{Ex436}$) for each ROI ($F_f$) was normalized to the initial fluorescence intensity ratio ($F_i$) of the same ROI. At least two different fields were sampled for each biological replicate, and the average emission response ($F_f/F_i$) with standard error of the mean is reported for all ROIs from three biological replicates (Figure S22 and Supplemental Video 5).

### FIGURES

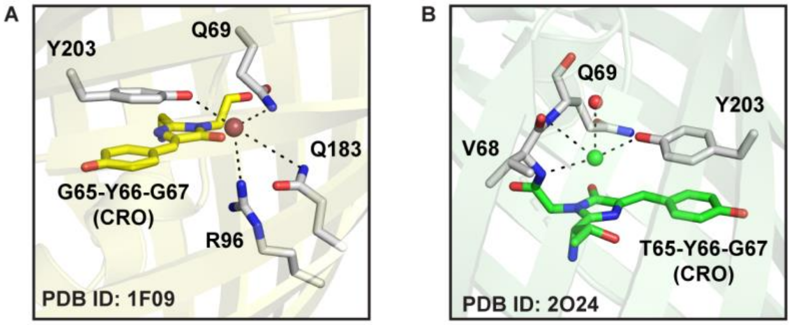


**Figure S1.** Representative anion-sensitive fluorescent proteins derived from the green fluorescent protein found in the jellyfish *Aequorea victoria* that have been structurally characterized with a bound anion. X-ray crystal structures of the binding pockets are shown for (A) avYFP-H148Q bound to iodide (purple sphere) and (B) $E^2$GFP bound to chloride (green sphere). The residues that make up the chromophore (CRO) are shown as colored sticks, and the anion coordinating residues within 4 Å are shown as grey sticks with dashed lines to the anion. Water molecules are shown as red spheres. All residues are labeled with the single letter amino acid abbreviation and corresponding position number.

atg ggc agc agc cat cat cat cat cat cac agc agc ggc ctg gtg
Met Gly Ser Ser His His His His His His Ser Ser Gly Leu Val

ccg cgc ggc agc **cat ATG** GTG AGC AAG GGC GAG GAA GAC AAC ATG
Pro Arg Gly Ser **His Met** Val Ser Lys Gly Glu Glu Asp Asn Met

GCG AGC CTG CCG GCG ACC CAT GAG CTG CAC ATC TTC GGC AGC ATT
1 Ala Ser Leu Pro Ala Thr His Glu Leu His Ile Phe Gly Ser Ile

AAC GGT GTG GAC TTT GAT ATG GTT GGT CAG GGC ACC GGT AAC CCG
16 Asn Gly Val Asp Phe Asp Met Val Gly Gln Gly Thr Gly Asn Pro

AAC GAC GGC TAC GAG GAA CTG AAC CTG AAG AGC ACC AAA GGT GAT
31 Asn Asp Gly Tyr Glu Glu Leu Asn Leu Lys Ser Thr Lys Gly Asp

CTG CAA TTC AGC CCG TGG ATT CTG GTG CCG CAC ATT GGC TAT GGT
46 Leu Gln Phe Ser Pro Trp Ile Leu Val Pro His Ile Gly Tyr Gly

TTT CAC CAG TAT CTG CCG TAT CCG GAT GGT ATG AGC CCG TTC CAA
61 Phe His Gln Tyr Leu Pro Tyr Pro Asp Gly Met Ser Pro Phe Gln

GCG GCG ATG GTG GAT GGC AGC GGT TAC CAG GTT CAC CGT ACC ATG
76 Ala Ala Met Val Asp Gly Ser Gly Tyr Gln Val His Arg Thr Met

CAA TTT GAA GAC GGT GCG AGC CTG ACC GTT AAC TAC CGT TAT ACC
91 Gln Phe Glu Asp Gly Ala Ser Leu Thr Val Asn Tyr Arg Tyr Thr

TAC GAG GGC AGC CAC ATC AAG GGT GAA GCG CAG GTG AAG GGT ACC
106 Tyr Glu Gly Ser His Ile Lys Gly Glu Ala Gln Val Lys Gly Thr

GGT TTC CCG GCG GAT GGT CCG GTT ATG ACC AAC AGC CTG ACC GCG
121 Gly Phe Pro Ala Asp Gly Pro Val Met Thr Asn Ser Leu Thr Ala

GCG GAC TGG TGC CGT AGC AAG **AAA** ACC TAT CCG AAC GAT AAG ACC
136 Ala Asp Trp Cys Arg Ser Lys **Lys** Thr Tyr Pro Asn Asp Lys Thr

ATC ATT AGC ACC TTT AAA TGG AGC TAT ACC ACC GGC AAC GGT AAA
151 Ile Ile Ser Thr Phe Lys Trp Ser Tyr Thr Thr Gly Asn Gly Lys

CGT TAC CGT AGC ACC GCG CGT ACC ACC TAT ACC TTT GCG AAG CCG
166 Arg Tyr Arg Ser Thr Ala Arg Thr Thr Tyr Thr Phe Ala Lys Pro

ATG GCG GCG AAC TAT CTG AAA AAC CAG CCG ATG TAC GTG TTC **CGT**
181 Met Ala Ala Asn Tyr Leu Lys Asn Gln Pro Met Tyr Val Phe **Arg**

AAG ACC GAG CTG AAG CAC AGC AAA ACC GAG CTG AAC TTC AAG GAA
196 Lys Thr Glu Leu Lys His Ser Lys Thr Glu Leu Asn Phe Lys Glu

TGG CAA AAA GCG TTT ACC GAC GTT ATG GGT ATG GAT GAA CTG TAC
211 Trp Gln Lys Ala Phe Thr Asp Val Met Gly Met Asp Glu Leu Tyr

AAA tga **gga tcc**
226 Lys * **Gly Ser**

**Figure S2.** The nucleotide (top row) and amino acid (bottom) sequences of the mNeonGreen (mNG) construct used to generate the double site-saturation mutagenesis library. The nucleotide sequence for mNG (green) (UniProt ID: A0A1S4NYF2) was cloned into the pET-28b(+) vector (black) between the NdeI and BamHI restriction sites (black and bold) with an N-terminal polyhistidine tag and a C-terminal stop codon (*). The K143 and R195 sites targeted for mutagenesis are bolded and highlighted in yellow. The mNG amino acid sequence numbering corresponds to the sequence reported for PDB ID: 5LTP.

**Table S1.** The primers used to generate the double site-saturation mutagenesis library at the K143 and R195 sites (shown in red) in mNG.

| Description | Primer Sequence (5' to 3') |
|---|---|
| K143 SSM Forward | GACTGGTGCCGTAGCAAGNDTACCTATCCGAACGATA |
| | GACTGGTGCCGTAGCAAGVHGACCTATCCGAACGATA |
| | GACTGGTGCCGTAGCAAGTGGACCTATCCGAACGATA |
| K143 SSM Reverse | CTTGCTACGGCACCAGTCCGCCGCGGTCAGGCTG |
| R195 SSM Forward | CAGCCGATGTACGTGTTCNDTAAGACCGAGCTGAAGC |
| | CAGCCGATGTACGTGTTCVHGAAGACCGAGCTGAAGC |
| | CAGCCGATGTACGTGTTCTGGAAGACCGAGCTGAAGC |
| R195 SSM Reverse | GAACACGTACATCGGCTGGTTTTTCAGATAGTTC |

**Table S2.** Polymerase chain reaction conditions to generate the double site-saturation mutagenesis library at K143 and R195.

<table>
<tr><th colspan="4">Reaction Conditions</th></tr>
<tr><th colspan="2">Solution</th><th>Concentration</th><th>Volume (µL)</th></tr>
<tr><td colspan="2">Template</td><td>10 ng/µL</td><td>5</td></tr>
<tr><td colspan="2">K143 SSM Forward Primer</td><td>10 µM</td><td>3</td></tr>
<tr><td colspan="2">K143 SSM Reverse Primer</td><td>10 µM</td><td>3</td></tr>
<tr><td colspan="2">R195 SSM Forward Primer</td><td>10 µM</td><td>3</td></tr>
<tr><td colspan="2">R195 SSM Reverse Primer</td><td>10 µM</td><td>3</td></tr>
<tr><td colspan="2">dNTPs</td><td>10 mM</td><td>1</td></tr>
<tr><td colspan="2">Phusion GC Buffer</td><td>5X</td><td>10</td></tr>
<tr><td colspan="2">Phusion DNA Polymerase</td><td>-</td><td>0.5</td></tr>
<tr><td colspan="2">Autoclaved Water</td><td>-</td><td>21.5</td></tr>
<tr><td colspan="2">Total</td><td>-</td><td>50</td></tr>
<tr><th colspan="4">Thermocycler Settings</th></tr>
<tr><th>Step</th><th>Temperature (° C)</th><th>Time (s)</th><th>Number of Cycles</th></tr>
<tr><td rowspan="2">Template Denaturation</td><td>95</td><td>30</td><td>1</td></tr>
<tr><td>95</td><td>10</td><td rowspan="3">30</td></tr>
<tr><td>Annealing</td><td>56.3</td><td>30</td></tr>
<tr><td>Short Extension</td><td>72</td><td>275</td></tr>
<tr><td>Long Extension</td><td>72</td><td>600</td><td>1</td></tr>
<tr><td>Storage</td><td>72</td><td>∞</td><td>1</td></tr>
</table>

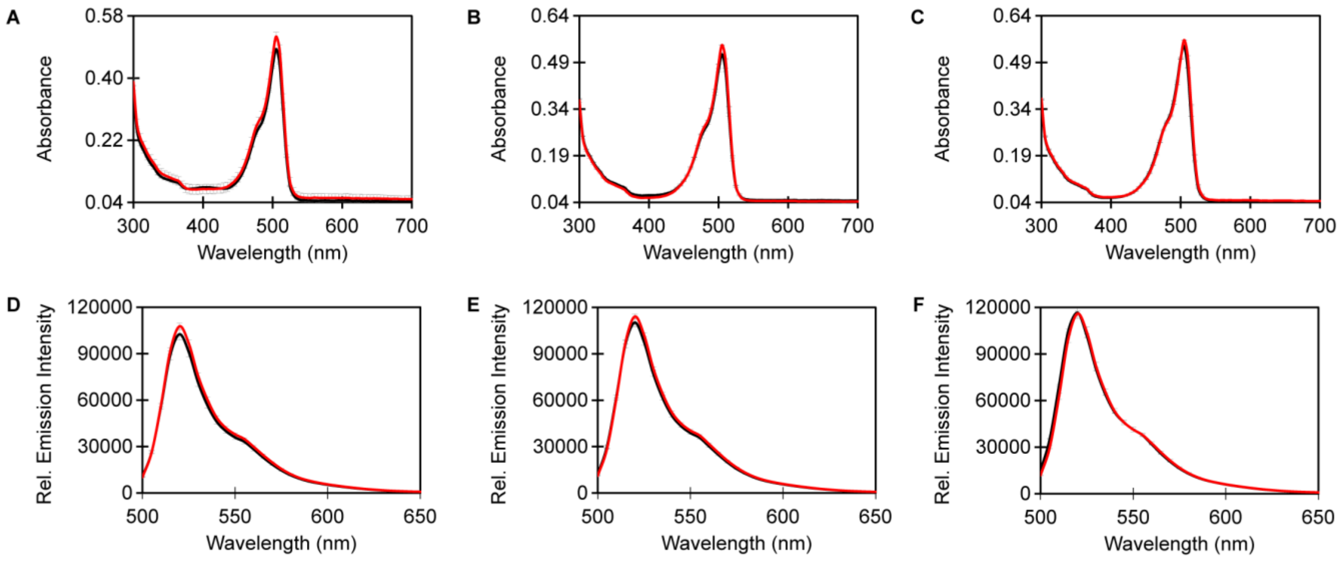


**Figure S3.** Absorption spectra of 10 µM mNG in the absence (black line) and presence (red) of 197 mM NaCl in 50 mM sodium phosphate buffer at (A) pH 6, (B) pH 7 and (C) pH 8. Emission spectra of mNG at (D) pH 6, (E) pH 7, and (F) pH 8 ($\lambda_{ex}$ = 485 nm, $\lambda_{em}$ = 500–650 nm). The average of two technical replicates from one protein preparation with standard deviation is reported.

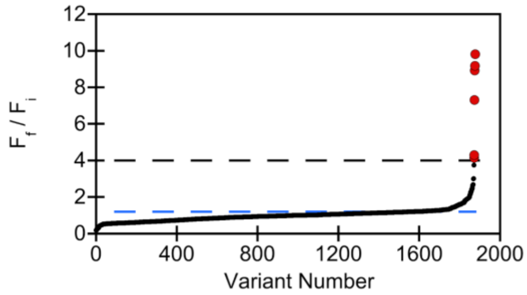


**Figure S4.** Summary plot of the double site-saturation mutagenesis library for the 1,880 variants (black circles) screened in the absence ($F_i$) and presence ($F_f$) of 100 mM NaBr in 25 mM sodium phosphate buffer at pH 8. The average turn-on fluorescence response ($F_f/F_i$) for eight biological replicates of the mNG parent is indicated by the blue dashed line ($F_f/F_i$ = 1.2-fold), and the threshold to identify improved variants is indicated by the black dashed line ($F_f/F_i$ > 4-fold). The six improved variants selected for rescreening are indicated with red circles (Table S3).

**Table S3.** Summary of the double site-saturation mutagenesis library screen and rescreening results from *E. coli* lysates. The turn-on fluorescence response ($F_f/F_i$) of mNG and the top six variants are shown.

| Protein | Codons at K143/R195 | Mutations | $F_f/F_i$ for $Br^-$ (pH 8)[a] | $F_f/F_i$ for $Cl^-$ (pH 7)[b] | $F_f/F_i$ for $Cl^-$ (pH 8)[b] |
|---|---|---|---|---|---|
| mNG | AAA/CGT | – | 1.2 | – | – |
| P8 A1 | GGT/AAG | K143G/R195K | 4.3 | 2.7 ± 2 | 2.8 ± 0.2 |
| P20 F12 | CTG/ATT | K143L/R195I | 4.2 | 4.5 ± 1 | 3.1 ± 0.2 |
| P3 B10 | TAT/ATT | K143Y/R195I | 9.8 | 11 ± 4 | 5.9 ± 0.3 |
| P11 H3 (ChlorON-1) | TGG/CTG | K143W/R195L | 7.3 | 29 ± 2 | 25 ± 1 |
| P6 C8 (ChlorON-2) | CGT/ATT | K143R/R195I | 8.9 | 25 ± 4 | 9 ± 2 |
| P5 A9 (ChlorON-3) | CGT/CTG | K143R/R195L | 9.2 | 30 ± 4 | 11 ± 2 |

[a]*Fluorescence response from the library screening with 100 mM NaBr in 25 mM sodium phosphate buffer at pH 8.*
[b]*Rescreening results with 100 mM NaCl in 25 mM sodium phosphate buffer at pH 7 and 8. The average of two biological replicates with standard error of the mean is reported.*

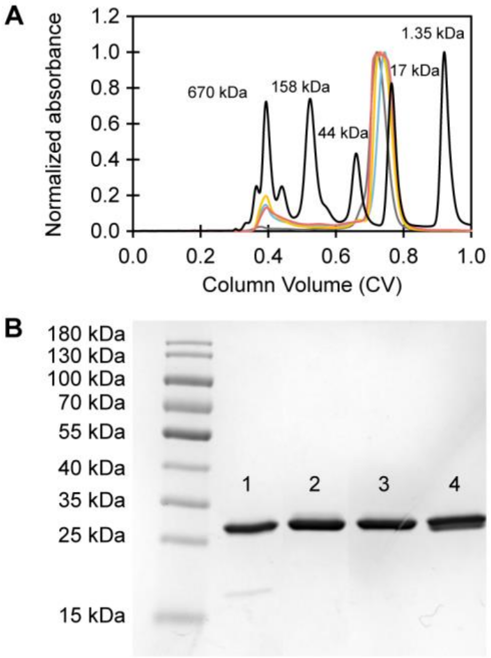


**Figure S5.** Analysis of the oligomeric state and purity for the ChlorON sensors and mNG. (A) Representative size-exclusion chromatographs for ChlorON-1 (red), ChlorON-2 (blue), ChlorON-3 (yellow), and mNG (gray) in 50 mM sodium phosphate buffer at pH 7.4 containing 200 mM NaCl. The thyroglobulin (670 kDa), $\gamma$-globulin (158 kDa), ovalbumin (44 kDa), myoglobulin (17 kDa), and vitamin B12 (1.35 kDa) protein standards are shown in black. For each protein, the normalized absorbance traces at 280 nm are shown. (B) Representative image of a Coomassie-stained SDS-PAGE for purified ChlorON-1 (1), ChlorON-2 (2), ChlorON-3 (3), and mNG (4). The theoretical molecular weight for each protein is ~26.6 kDa.

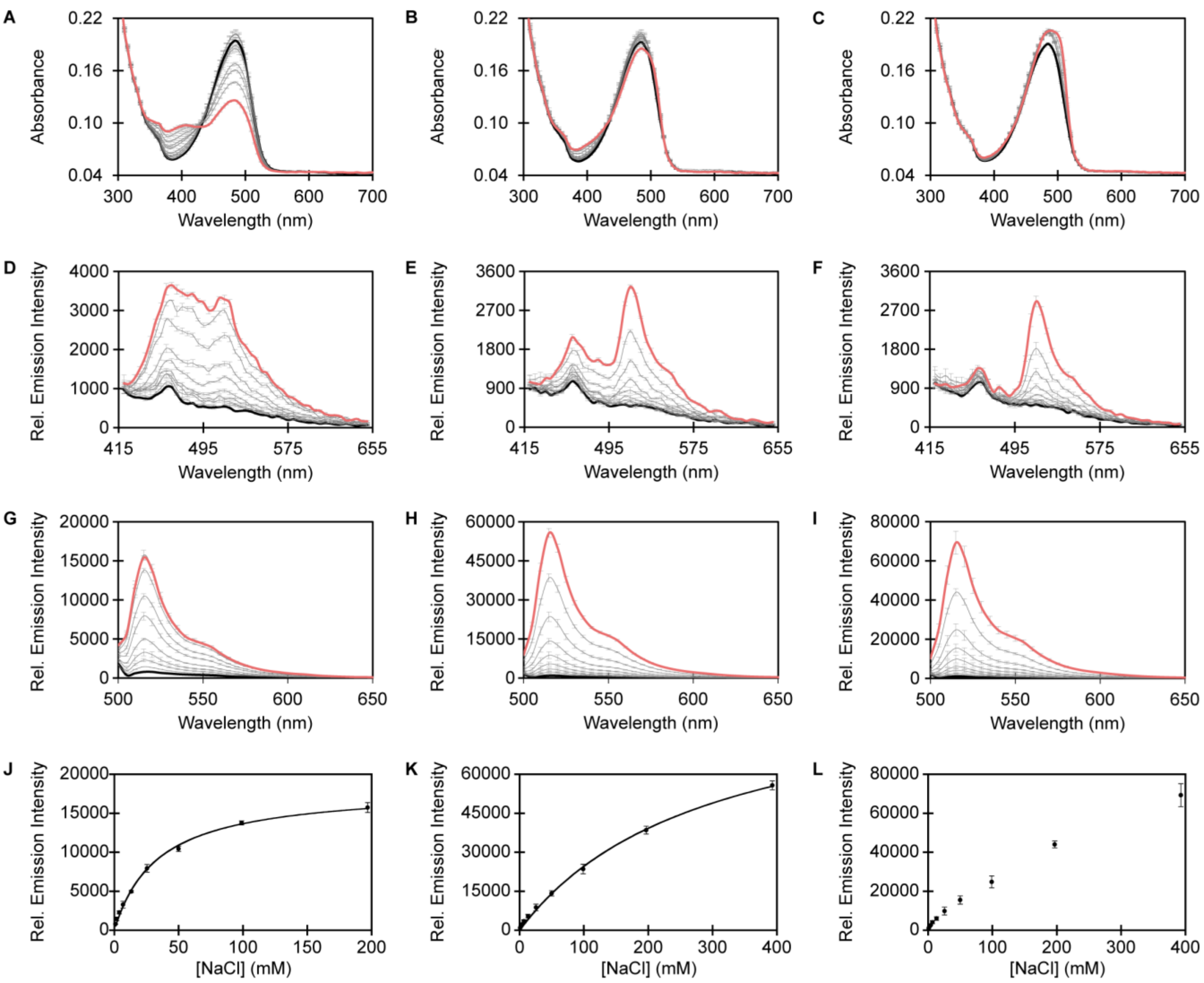


**Figure S6.** Spectroscopic characterization of ~10 µM ChlorON-1 in the presence of 1 (bold black), 2, 3.9, 6.9, 13.3, 25.5, 50, 99, 197, and 393 mM (red) NaCl. Absorption spectra of ChlorON-1 in 50 mM sodium phosphate buffer at (A) pH 6, (B) pH 7, and (C) pH 8. Emission spectra of ChlorON-1 at (D) pH 6, (E) pH 7, and (F) pH 8 ($\lambda_{ex}$ = 400 nm, $\lambda_{em}$ = 415–650 nm). Emission spectra of ChlorON-1 at (G) pH 6, (H) pH 7, and (I) pH 8 ($\lambda_{ex}$ = 485 nm, $\lambda_{em}$ = 500–650 nm). The relative emission responses ($\lambda_{ex}$ = 485 nm, $\lambda_{em}$ = 515 nm) from panels G–I were used to determine the apparent dissociation constant for chloride binding to ChlorON-1 at (J) pH 6 ($K_d$ = 39 ± 5 mM), (K) pH 7 ($K_d$ = 285 ± 59 mM), and (L) pH 8 ($K_d$ = not determined, $R^2$ = 0.98). The average of four technical replicates from two protein preparations with standard error of the mean is reported (Table S4).

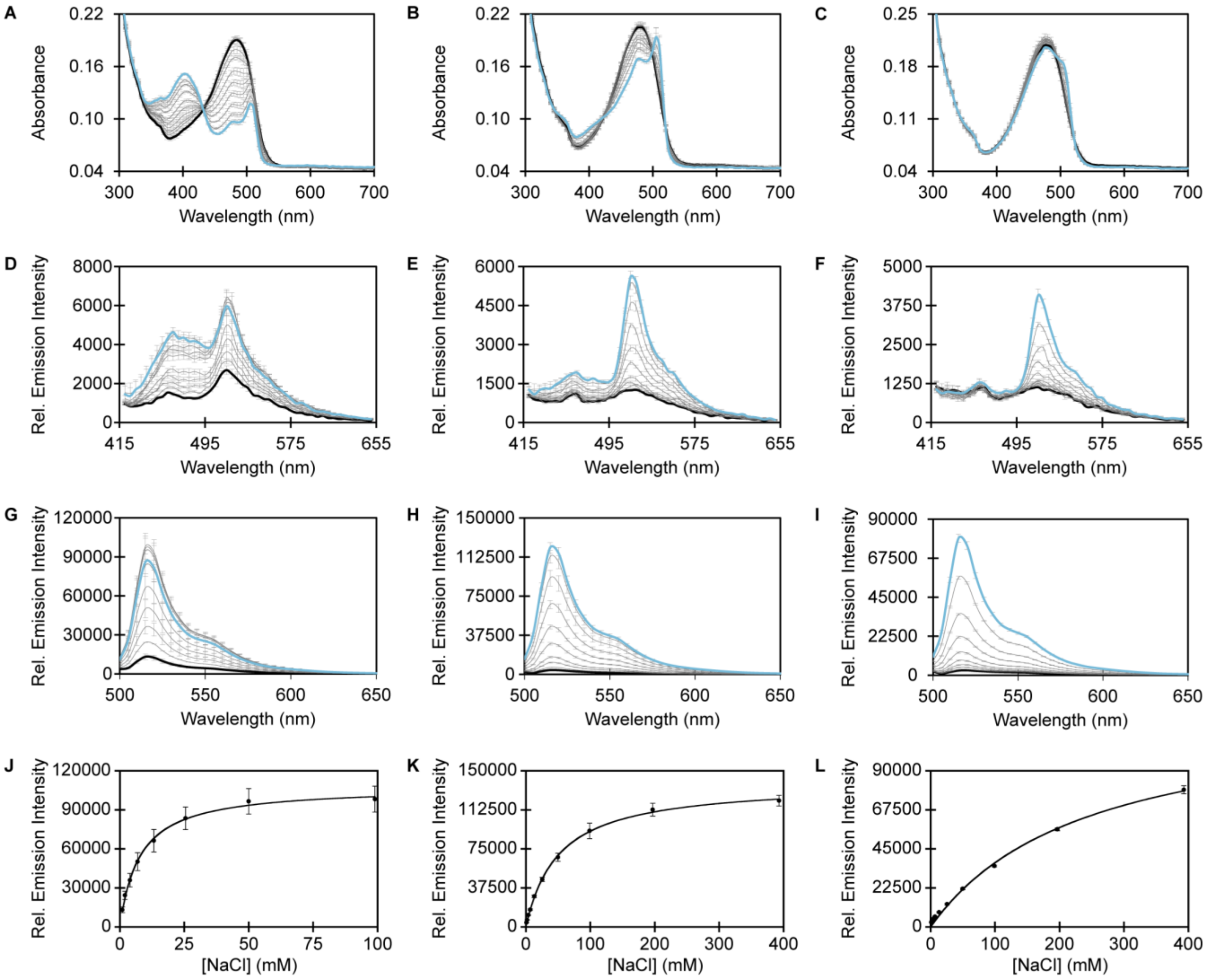


**Figure S7.** Spectroscopic characterization of ~10 µM ChlorON-2 in the presence of 1 (bold black), 2, 3.9, 6.9, 13.3, 25.5, 50, 99, 197, and 393 mM (blue) NaCl. Absorption spectra of ChlorON-2 in 50 mM sodium phosphate buffer at (A) pH 6, (B) pH 7, and (C) pH 8. Emission spectra of ChlorON-2 at (D) pH 6, (E) pH 7, and (F) pH 8 ($\lambda_{ex}$ = 400 nm, $\lambda_{em}$ = 415–650 nm). Emission spectra of ChlorON-2 at (G) pH 6, (H) pH 7, and (I) pH 8 ($\lambda_{ex}$ = 485 nm, $\lambda_{em}$ = 500–650 nm). The relative emission responses ($\lambda_{ex}$ = 485 nm, $\lambda_{em}$ = 515 nm) from panels G–I were used to determine the apparent dissociation constant for chloride binding to ChlorON-2 at (J) pH 6 ($K_d$ = 7.5 ± 1 mM), (K) pH 7 ($K_d$ = 55 ± 5 mM), and (L) pH 8 ($K_d$ = 228 ± 25). The average of four technical replicates from two protein preparations with standard error of the mean is reported (Table S4).

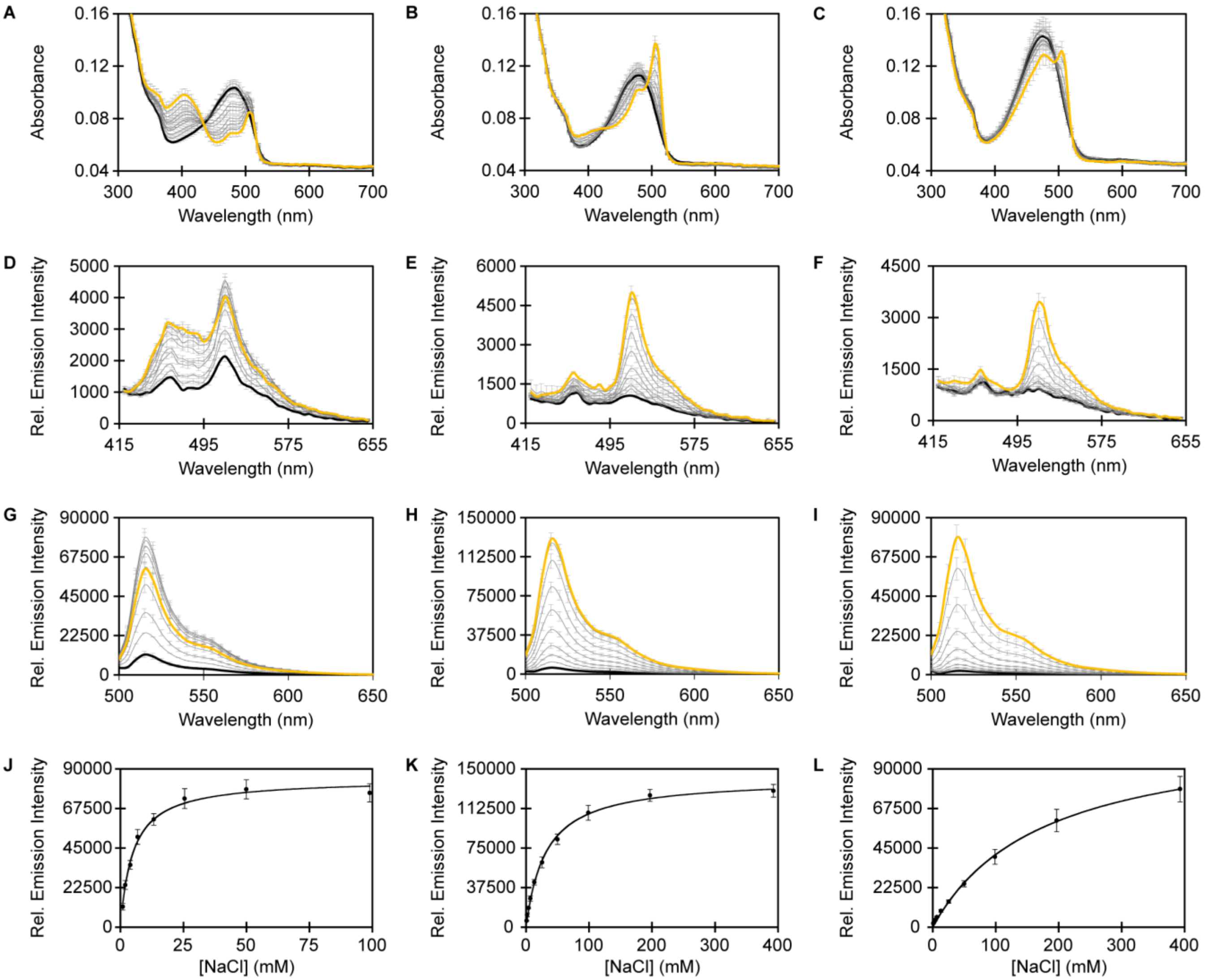


**Figure S8.** Spectroscopic characterization of ~10 µM ChlorON-3 in the presence of 1 (bold black), 2, 3.9, 6.9, 13.3, 25.5, 50, 99, 197, and 393 mM (yellow) NaCl. Absorption spectra of ChlorON-3 in 50 mM sodium phosphate buffer at (A) pH 6, (B) pH 7, and (C) pH 8. Emission spectra of ChlorON-3 at (D) pH 6, (E) pH 7, and (F) pH 8 ($\lambda_{ex}$ = 400 nm, $\lambda_{em}$ = 415–650 nm). Emission spectra of ChlorON-3 at (G) pH 6, (H) pH 7, and (I) pH 8 ($\lambda_{ex}$ = 485 nm, $\lambda_{em}$ = 500–650 nm). The relative emission responses ($\lambda_{ex}$ = 485 nm, $\lambda_{em}$ = 515 nm) from panels G–I were used to determine the apparent dissociation constant for chloride binding to ChlorON-3 at (J) pH 6 ($K_d$ = 4.4 ± 0.9 mM), (K) pH 7 ($K_d$ = 30 ± 1 mM), and (L) pH 8 ($K_d$ = 169 ± 79). The average of four technical replicates from two protein preparations with standard error of the mean is reported (Table S4).

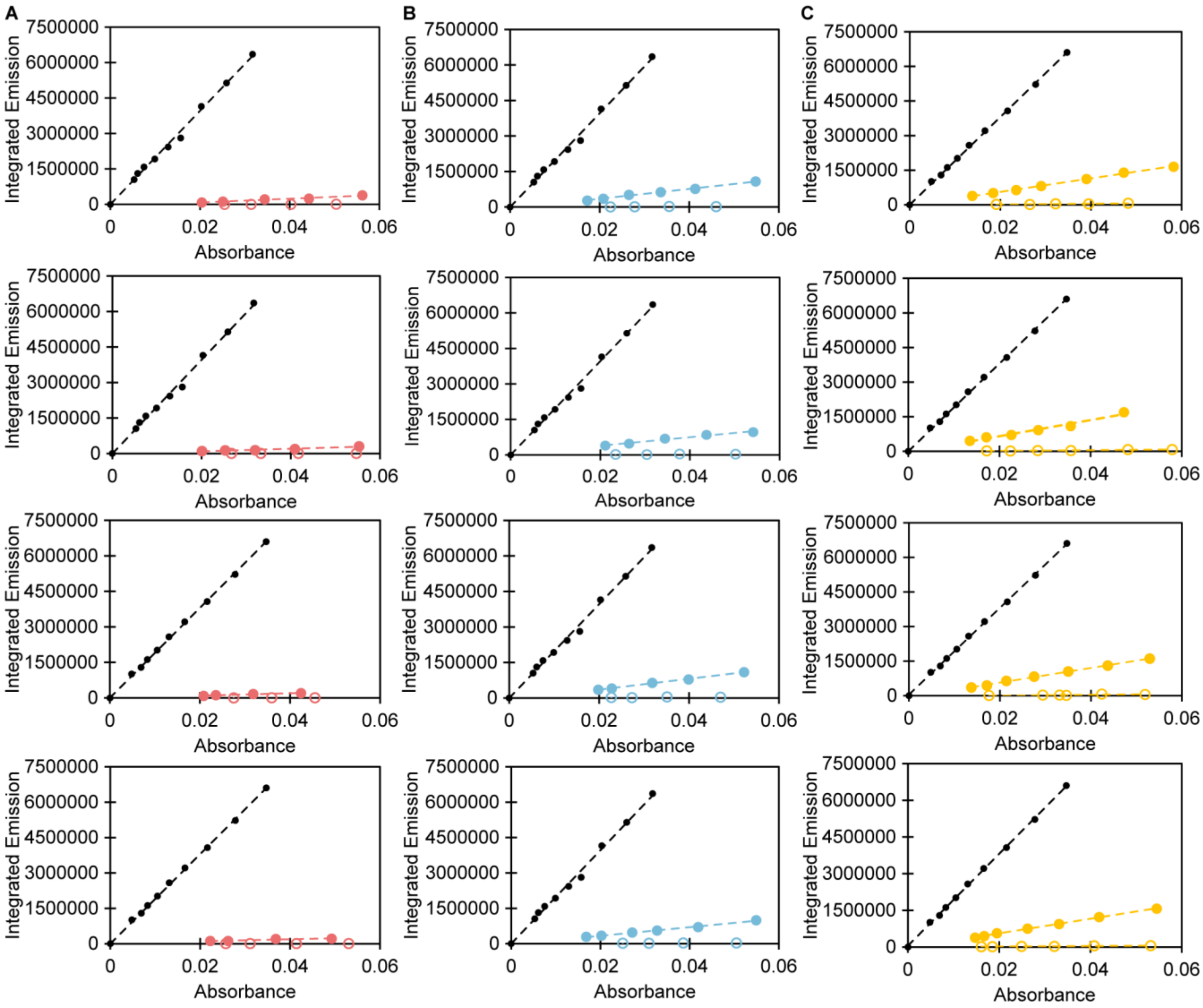


**Figure S9.** Fluorescence quantum yield curves of (A) ChlorON-1, (B) ChlorON-2, and (C) ChlorON-3 in presence of 0 mM (open) and 200 mM (filled circles) NaCl in 50 mM sodium phosphate buffer at pH 7. For each sensor, four technical replicates from two protein preparations are shown along with the fluorescein standard curves in 100 mM NaOH (black circles). In each graph, the integrated emission ($\lambda_{ex}$ = 488 nm, $\lambda_{em}$ = 490–750 nm) is plotted versus the corresponding absorbance intensity at 488 nm ($R^2 > 0.99$). The slope of each graph was determined to calculate the quantum yields.

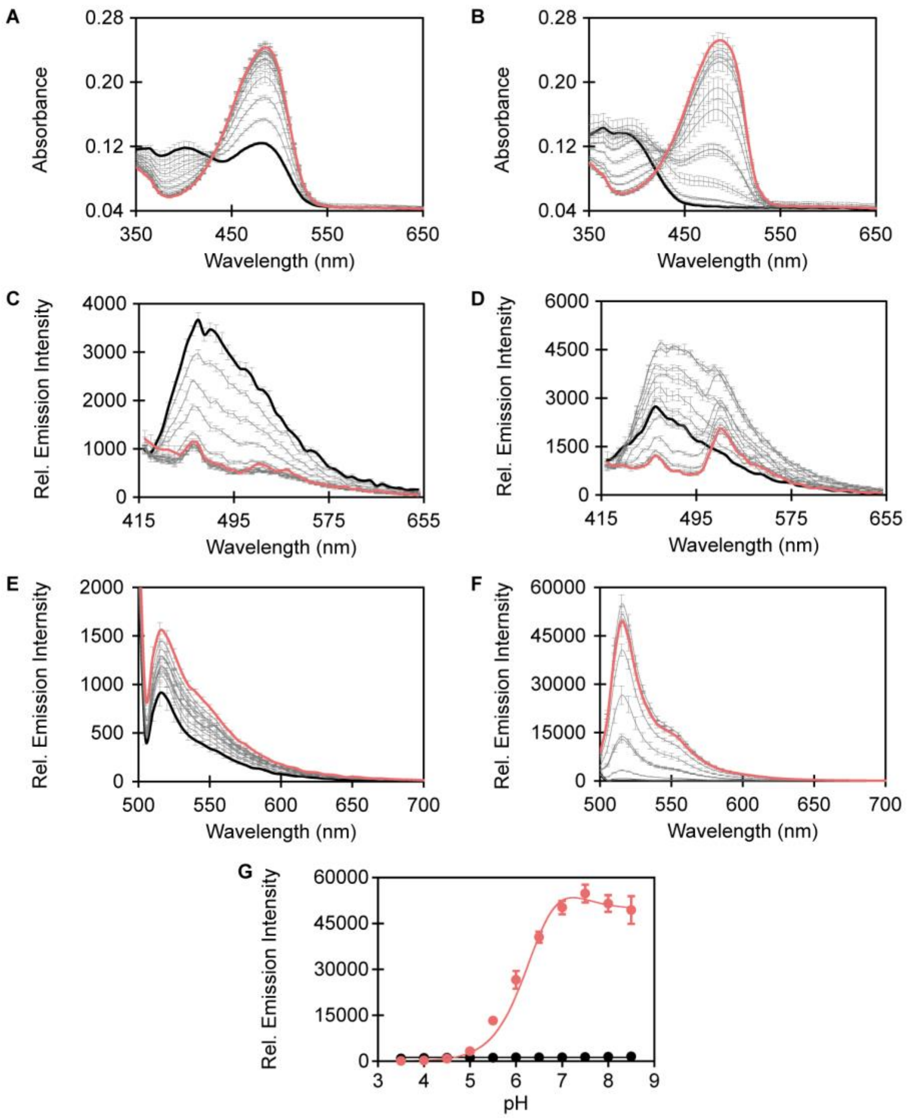


**Figure S10.** Spectroscopic characterization of ~10 µM ChlorON-1 in 50 mM sodium acetate buffer from pH 3.5 (black)–5.5 and 50 mM sodium phosphate buffer from pH 5.5–8.5 (red). Absorption spectra of ChlorON-1 in the presence of (A) 1 mM and (B) 197 mM NaCl. Emission spectra of ChlorON-1 in the presence of (C) 1 mM and (D) 197 mM NaCl ($\lambda_{ex}$ = 400 nm, $\lambda_{em}$ = 415–650 nm). Emission spectra of ChlorON-1 in the presence of (E) 1 mM and (F) 197 mM NaCl ($\lambda_{ex}$ = 485 nm, $\lambda_{em}$ = 500–700 nm). (G) The relative emission responses ($\lambda_{ex}$ = 485 nm, $\lambda_{em}$ = 515 nm) from panels E and F were used to determine the p$K_a$s of ChlorON-1 in the presence of 1 mM (black circles, p$K_a$ = not determined) and 197 mM NaCl (red circles, p$K_a$ = 6.0 ± 0.1). The average of four technical replicates from two protein preparations with standard error of the mean is reported (Table S4).

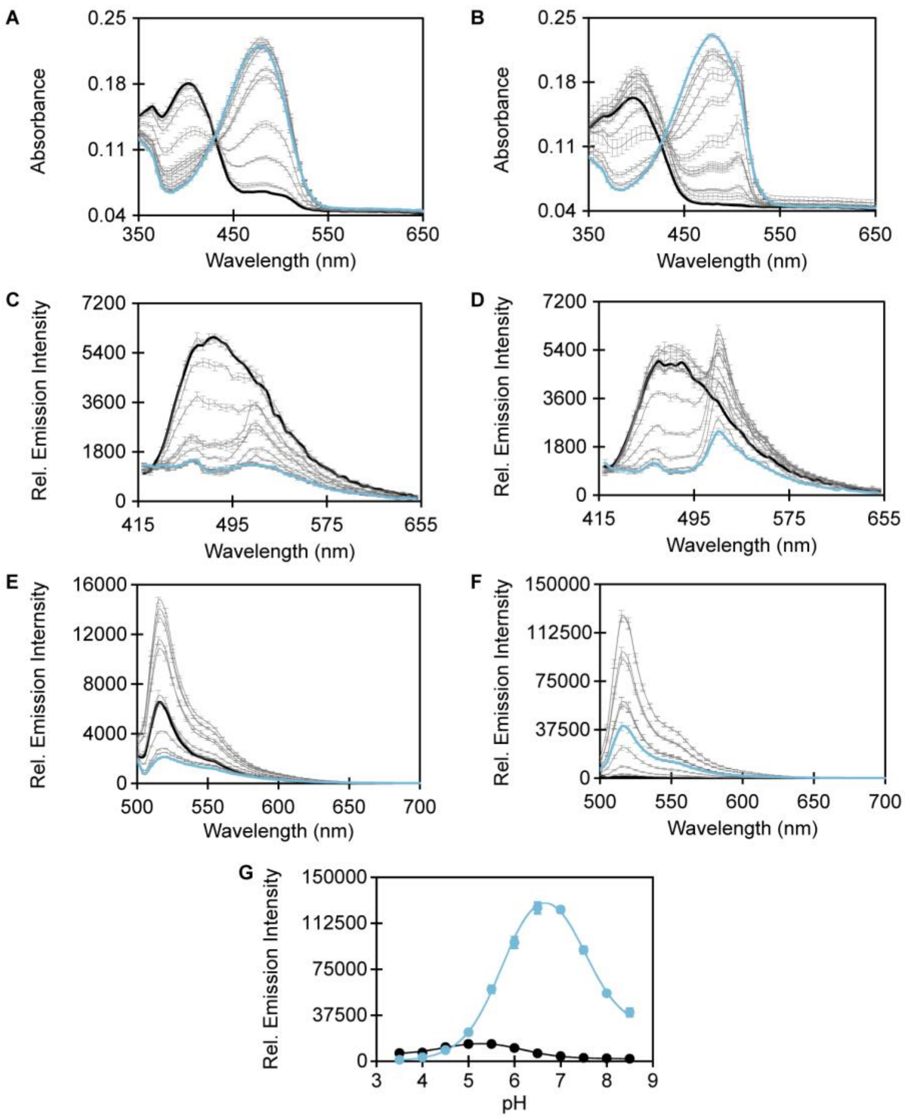


**Figure S11.** Spectroscopic characterization of ~10 µM ChlorON-2 in 50 mM sodium acetate buffer from pH 3.5 (black)–5.5 and 50 mM sodium phosphate buffer from pH 5.5–8.5 (blue). Absorption spectra of ChlorON-2 in the presence of (A) 1 mM and (B) 197 mM NaCl. Emission spectra of ChlorON-2 in the presence of (C) 1 mM and (D) 197 mM NaCl ($\lambda_{ex}$ = 400 nm, $\lambda_{em}$ = 415–650 nm). Emission spectra of ChlorON-2 in the presence of (E) 1 mM and (F) 197 mM NaCl ($\lambda_{ex}$ = 485 nm, $\lambda_{em}$ = 500–700 nm). (G) The relative emission responses ($\lambda_{ex}$ = 485 nm, $\lambda_{em}$ = 515 nm) from panels E and F were used to determine the p$K_a$s of ChlorON-2 in the presence of 1 mM (black circles, p$K_a$1 = 4.6 ± 0.1, p$K_a$2 = 6.1 ± 0.1) and 197 mM NaCl (blue circles, p$K_a$1 = 5.7 ± 0.1, p$K_a$2 = 7.6 ± 0.1). The average of four technical replicates from two protein preparations with standard error of the mean is reported (Table S4).

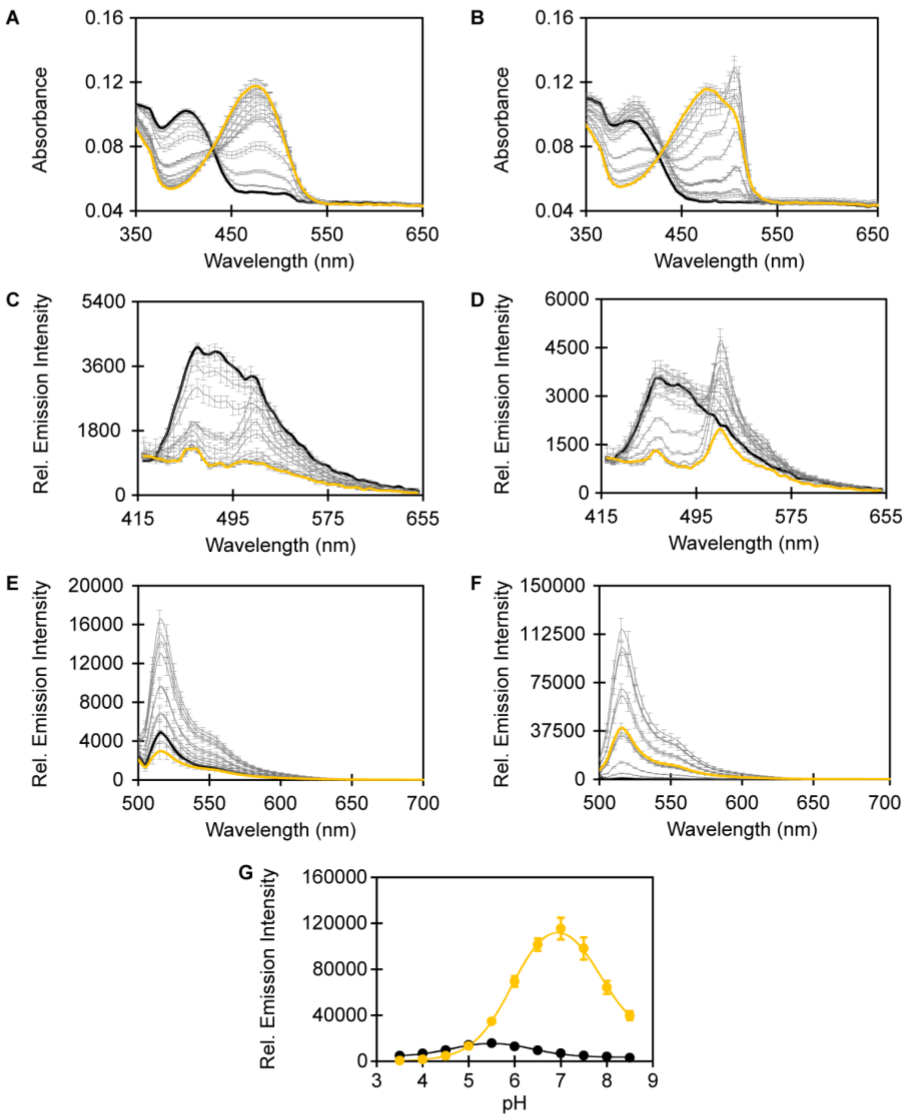


**Figure S12.** Spectroscopic characterization of ~10 µM ChlorON-3 in 50 mM sodium acetate buffer from pH 3.5 (black)–5.5 and 50 mM sodium phosphate buffer from pH 5.5–8.5 (yellow). Absorption spectra of ChlorON-3 in the presence of (A) 1 mM and (B) 197 mM NaCl. Emission spectra of ChlorON-3 in the presence of (C) 1 mM and (D) 197 mM NaCl ($\lambda_{ex}$ = 400 nm, $\lambda_{em}$ = 415–650 nm). Emission spectra of ChlorON-3 in the presence of (E) 1 mM and (F) 197 mM NaCl ($\lambda_{ex}$ = 485 nm, $\lambda_{em}$ = 500–700 nm). (G) The relative emission responses ($\lambda_{ex}$ = 485 nm, $\lambda_{em}$ = 515 nm) from panels E and F were used to determine the p$K_a$s of ChlorON-3 in the presence of 1 mM (black circles, p$K_a$1 = 4.8 ± 0.2, p$K_a$2 = 6.1 ± 0.2) and 197 mM NaCl (yellow circles, p$K_a$1 = 6.0 ± 0.01, p$K_a$2 = 7.9 ± 0.1). The average of four technical replicates from two protein preparations with standard error of the mean is reported (Table S4).

**Table S4.** Summary of the turn-on fluorescence responses ($F_f/F_i$), apparent dissociation constant ($K_d$), p$K_a$, extinction coefficients ($\varepsilon$), quantum yields ($\Phi$), and molar brightness ($\varepsilon$ x $\Phi$) for ChlorON-1, ChlorON-2, and ChlorON-3 with 197 mM NaCl. The average of four technical replicates from two protein preparations with the standard error of the mean is reported.

| **Protein** | **ChlorON-1** | **ChlorON-2** | **ChlorON-3** |
|---|---|---|---|
| **Response ($F_f/F_i$)** | | | |
| pH 6 | 20 ± 3 | 7.4 ± 1 | 6.2 ± 1 |
| pH 7 | 45 ± 3 | 27 ± 5 | 20 ± 1 |
| pH 8 | 44 ± 4 | 21 ± 2 | 29 ± 1 |
| **Apparent dissociation constant ($K_d$, mM)** | | | |
| pH 6 | 39 ± 5 | 7.5 ± 1 | 4.4 ± 0.9 |
| pH 7 | 285 ± 59 | 55 ± 5 | 30 ± 1 |
| pH 8 | — | 228 ± 25 | 169 ± 79 |
| **p$K_a$** | | | |
| p$K_a$ -$Cl^-$ | — | 4.6 ± 0.1, 6.1 ± 0.1 | 4.8 ± 0.2, 6.1 ± 0.2 |
| p$K_a$ +$Cl^-$ | 6.0 ± 0.1 | 5.7 ± 0.1, 7.6 ± 0.1 | 6.0 ± 0.01, 7.9 ± 0.1 |
| **Absorption and Emission Maximum (nm) at pH 7** | | | |
| -$Cl^-$ | 480 | 480 | 480 |
| +$Cl^-$ | 480 | 480, 505 | 480, 505 |
| **Extinction coefficient (x $10^3$ ($M^{-1}$ * $cm^{-1}$) at pH 7** | | | |
| $\varepsilon$ -$Cl^-$ at 480 nm | 27.4 ± 4 | 21.9 ± 1 | 15.6 ± 0.5 |
| $\varepsilon$ +$Cl^-$ at 480 nm | 24.2 ± 3 | 16.6 ± 2 | 12.5 ± 0.6 |
| $\varepsilon$ +$Cl^-$ at 506 nm | — | 20.9 ± 4 | 18.7 ± 0.8 |
| **Quantum yield at pH 7** | | | |
| $\Phi$ -$Cl^-$ | < 0.01 | < 0.01 | < 0.01 |
| $\Phi$ +$Cl^-$ | 0.027 ± 0.007 | 0.093 ± 0.01 | 0.152 ± 0.01 |
| **Molar brightness at pH 7** | | | |
| -$Cl^-$ at 480 nm | 0.01 | 0.06 | 0.12 |
| +$Cl^-$ at 480 nm | 0.65 | 1.54 | 1.90 |
| +$Cl^-$ at 506 nm | — | 1.94 | 2.85 |

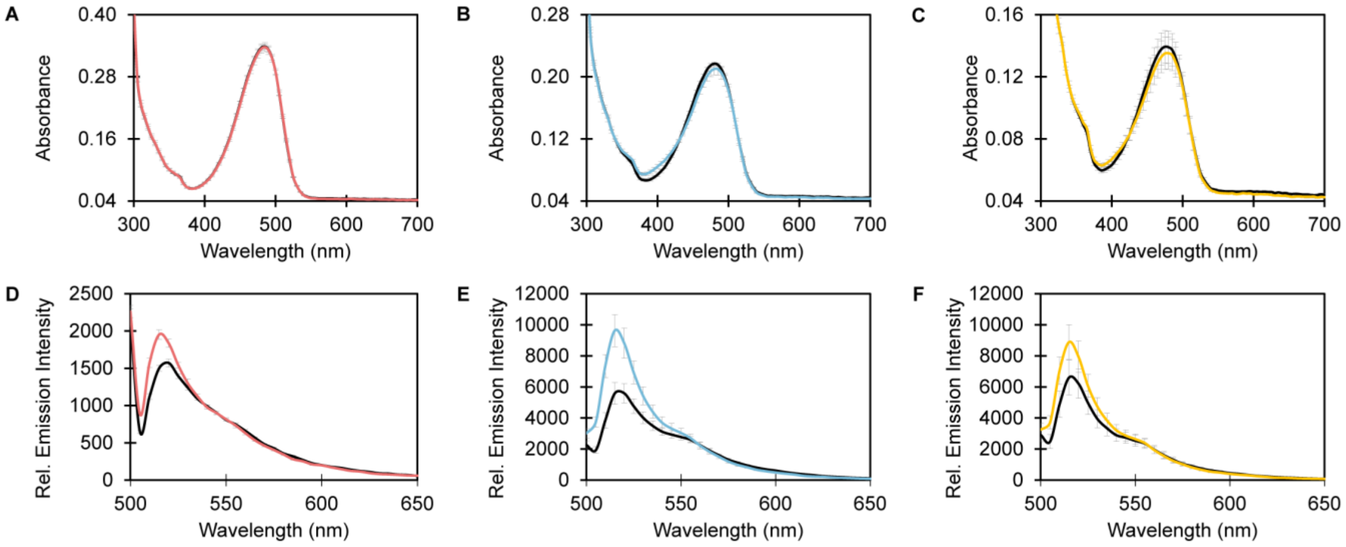


**Figure S13.** Spectroscopic characterization of ~10 µM ChlorON-1, ChlorON-2, and ChlorON-3 in the presence of 0 (bold black) and 196 mM (red, blue, and yellow, respectively) sodium phosphate in 50 mM HEPES buffer at pH 7. Absorption spectra of (A) ChlorON-1, (B) ChlorON-2, and (C) ChlorON-3. Emission spectra of (D) ChlorON-1, (E) ChlorON-2, and (F) ChlorON-3 ($\lambda_{ex}$ = 485 nm, $\lambda_{em}$ = 500–650 nm). The average of four technical replicates from two protein preparations with standard error of the mean is reported (Table S5).

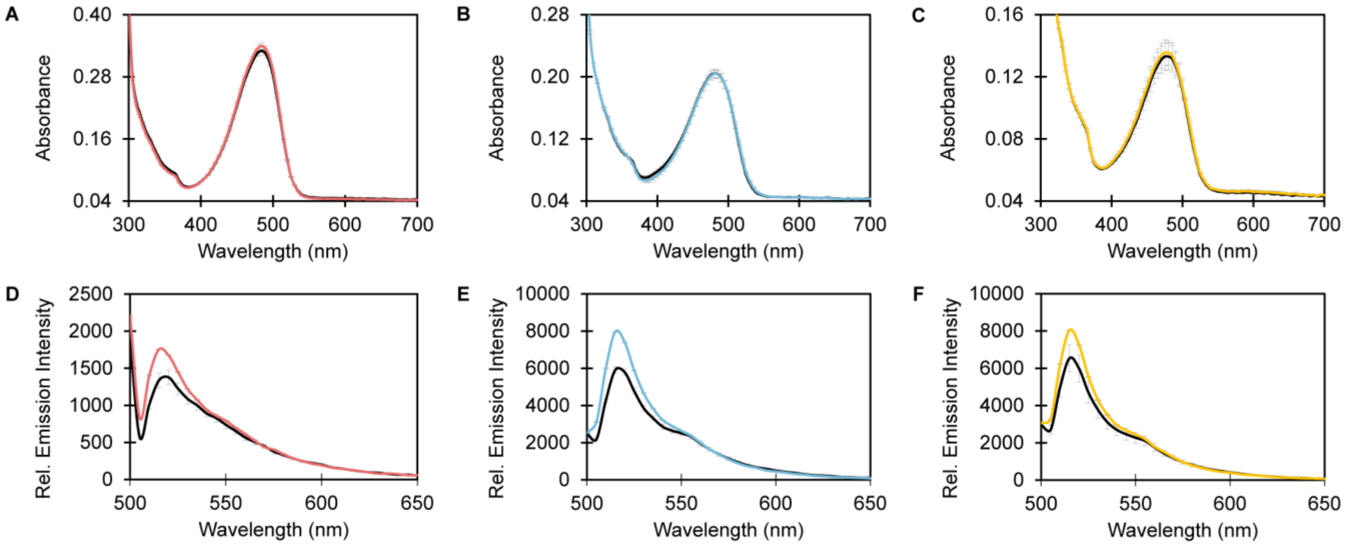


**Figure S14.** Spectroscopic characterization of ~10 µM ChlorON-1, ChlorON-2, and ChlorON-3 in the presence of 0 (bold black) and 196 mM (red, blue, and yellow, respectively) sodium acetate in 50 mM sodium phosphate buffer at pH 7. Absorption spectra of (A) ChlorON-1, (B) ChlorON-2, and (C) ChlorON-3. Emission spectra of (D) ChlorON-1, (E) ChlorON-2, and (F) ChlorON-3 ($\lambda_{ex}$ = 485 nm, $\lambda_{em}$ = 500–650 nm). The average of four technical replicates from two protein preparations with standard error of the mean is reported (Table S5).

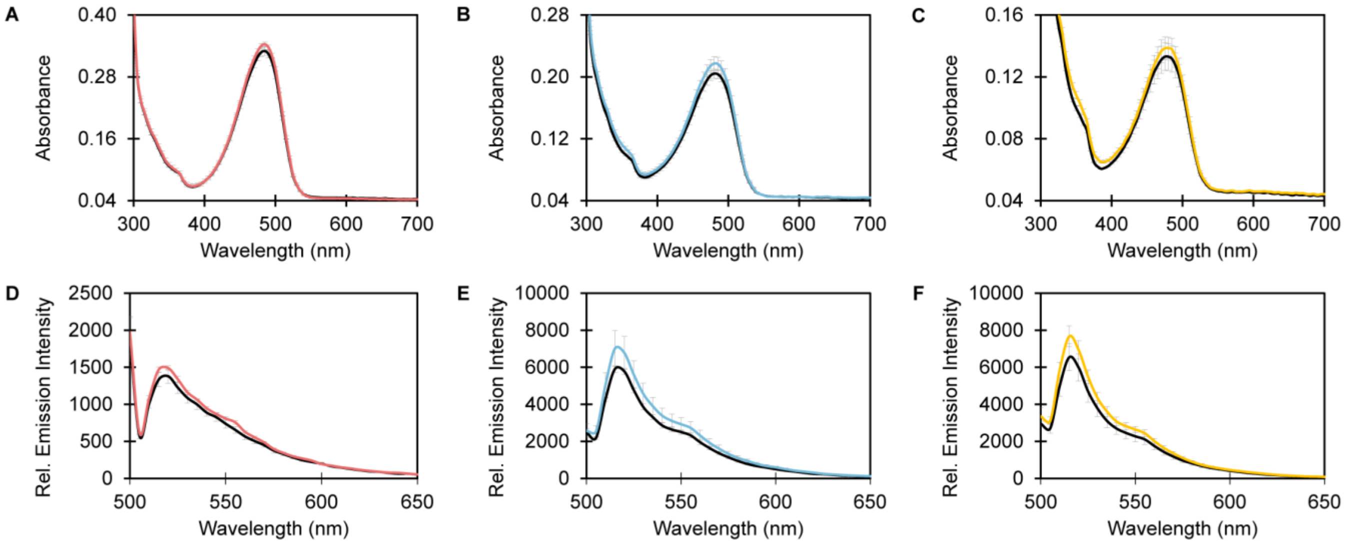


**Figure S15.** Spectroscopic characterization of ~10 µM ChlorON-1, ChlorON-2, and ChlorON-3 in the presence of 0 (bold black) and 196 mM (red, blue, and yellow, respectively) sodium citrate in 50 mM sodium phosphate buffer at pH 7. Absorption spectra of (A) ChlorON-1, (B) ChlorON-2, and (C) ChlorON-3. Emission spectra of (D) ChlorON-1, (E) ChlorON-2, and (F) ChlorON-3 ($\lambda_{ex}$ = 485 nm, $\lambda_{em}$ = 500–650 nm). The average of four technical replicates from two protein preparations with standard error of the mean is reported (Table S5).

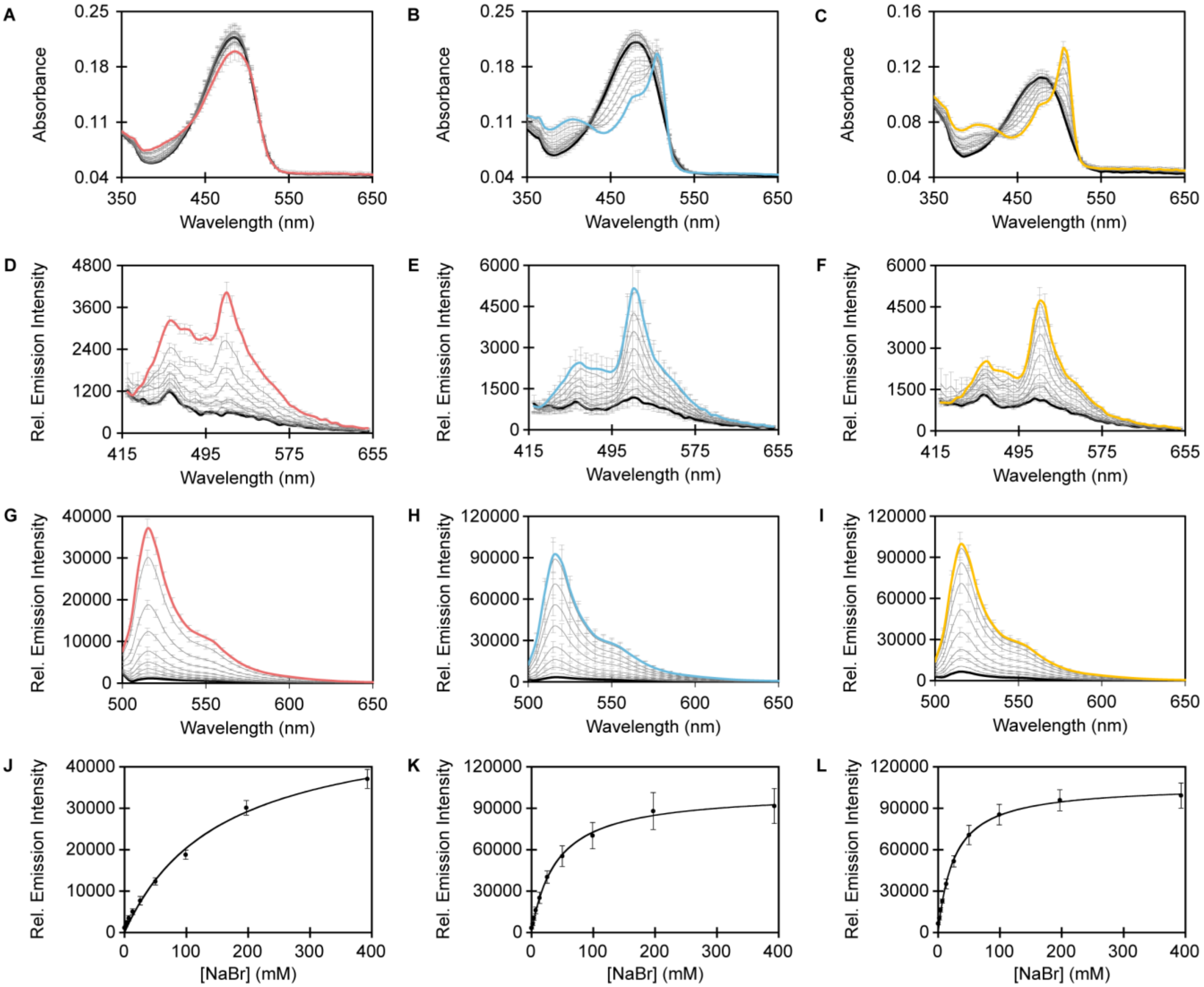


**Figure S16.** Spectroscopic characterization of ~10 µM ChlorON-1, ChlorON-2, and ChlorON-3 in the presence of 0 (bold black), 1.0, 2.9, 5.9, 12.3, 24.5, 49, 98, 196, and 392 mM (red, blue, and yellow, respectively) NaBr in 50 mM sodium phosphate buffer at pH 7. Absorption spectra of (A) ChlorON-1, (B) ChlorON-2, and (C) ChlorON-3. Emission spectra of (D) ChlorON-1, (E) ChlorON-2, and (F) ChlorON-3 ($\lambda_{ex}$ = 400 nm, $\lambda_{em}$ = 415–650 nm). Emission spectra of (G) ChlorON-1, (H) ChlorON-2, and (I) ChlorON-3 ($\lambda_{ex}$ = 485 nm, $\lambda_{em}$ = 500–650 nm). The relative emission responses ($\lambda_{ex}$ = 485 nm, $\lambda_{em}$ = 515 nm) from panels G–I were used to determine the apparent dissociation constants of bromide binding to (J) ChlorON-1 ($K_d$ = 204 ± 38), (K) ChlorON-2 ($K_d$ = 43 ± 1), and (L) ChlorON-3 ($K_d$ = 32 ± 2). The average of four technical replicates from two protein preparations with standard error of the mean is reported (Table S5).

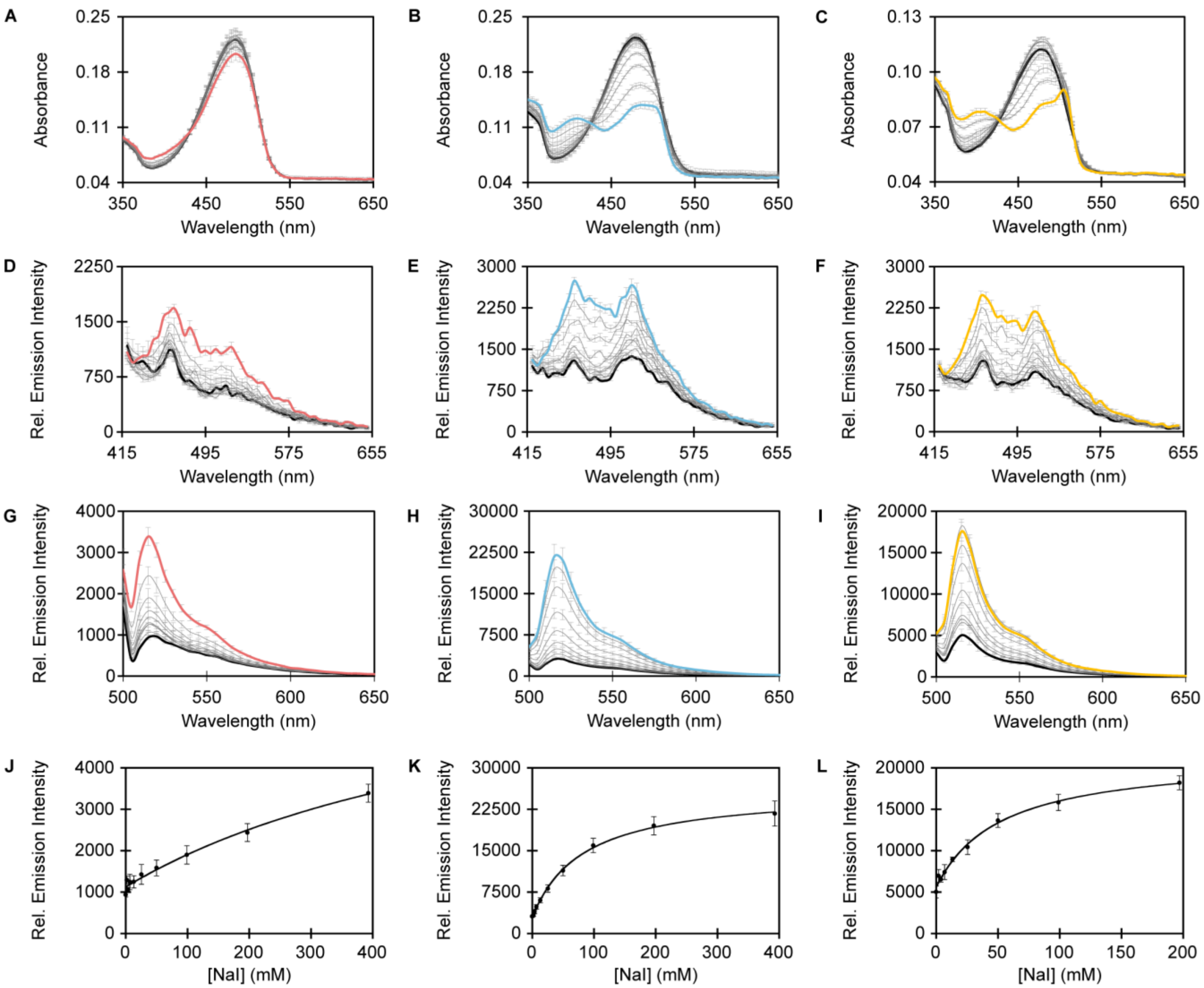


**Figure S17.** Spectroscopic characterization of ~10 µM ChlorON-1, ChlorON-2, and ChlorON-3 in the presence of 0 (bold black), 1.0, 2.9, 5.9, 12.3, 24.5, 49, 98, 196, and 392 mM (red, blue, and yellow, respectively) NaI in 50 mM sodium phosphate buffer at pH 7. Absorption spectra of (A) ChlorON-1, (B) ChlorON-2, and (C) ChlorON-3. Emission spectra of (D) ChlorON-1, (E) ChlorON-2, and (F) ChlorON-3 ($\lambda_{ex}$ = 400 nm, $\lambda_{em}$ = 415–650 nm). Emission spectra of (G) ChlorON-1, (H) ChlorON-2, and (I) ChlorON-3 ($\lambda_{ex}$ = 485 nm, $\lambda_{em}$ = 500–650 nm). The relative emission responses ($\lambda_{ex}$ = 485 nm, $\lambda_{em}$ = 515 nm) from panels G–I were used to determine the apparent dissociation constants of iodide binding to (J) ChlorON-1 ($K_d$ not determined, $R^2$ = 0.97), (K) ChlorON-2 ($K_d$ = 86 ± 4), and (L) ChlorON-3 ($K_d$ = 48 ± 7). The average of four technical replicates from two protein preparations with standard error of the mean is reported (Table S5).

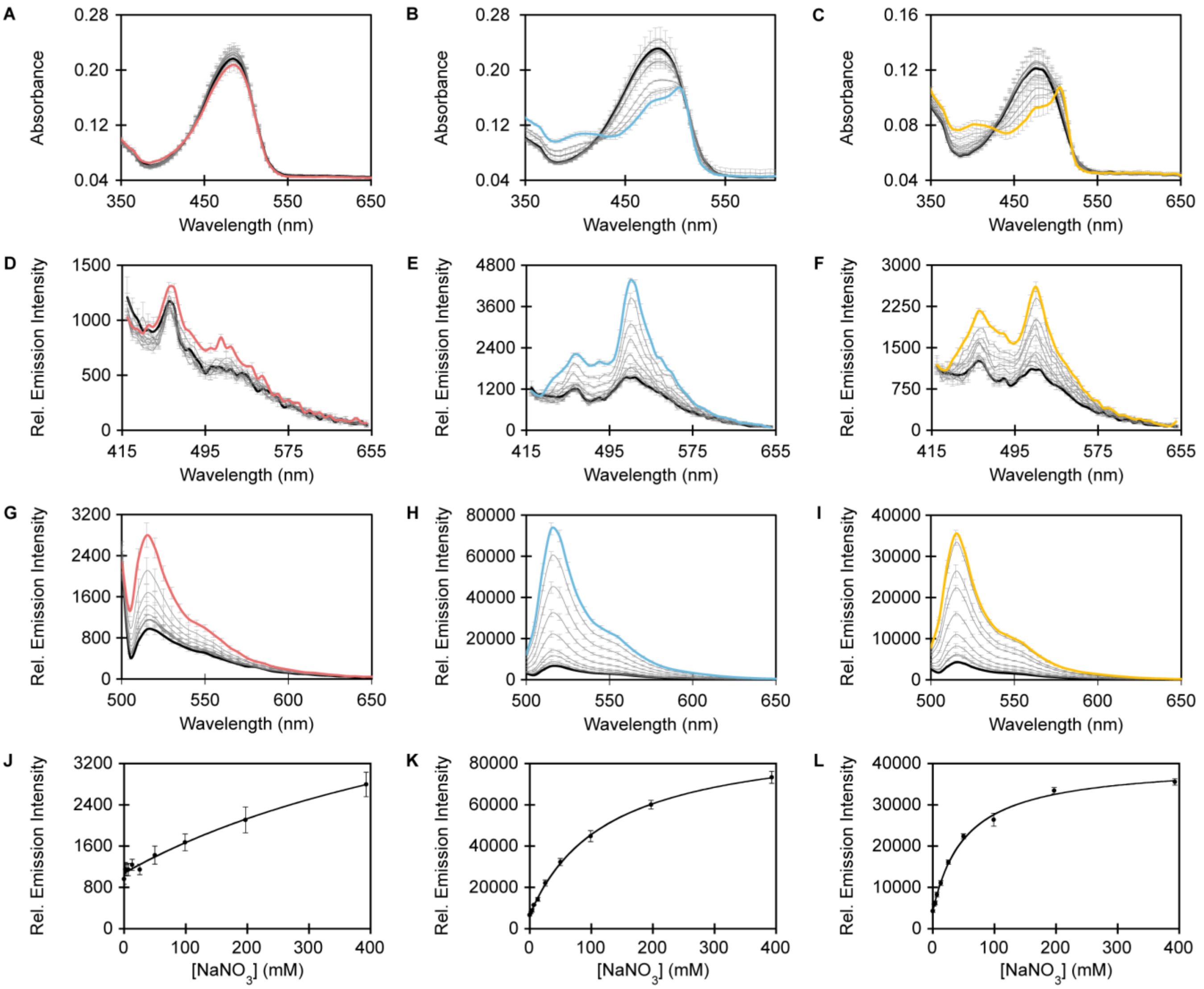


**Figure S18.** Spectroscopic characterization of ~10 µM ChlorON-1, ChlorON-2, and ChlorON-3 in the presence of 0 (bold black), 1.0, 2.9, 5.9, 12.3, 24.5, 49, 98, 196, and 392 mM (red, blue, and yellow, respectively) $NaNO_3$ in 50 mM sodium phosphate buffer at pH 7. Absorption spectra of (A) ChlorON-1, (B) ChlorON-2, and (C) ChlorON-3. Emission spectra of (D) ChlorON-1, (E) ChlorON-2, and (F) ChlorON-3 ($\lambda_{ex}$ = 400 nm, $\lambda_{em}$ = 415–650 nm). Emission spectra of (G) ChlorON-1, (H) ChlorON-2, and (I) ChlorON-3 ($\lambda_{ex}$ = 485 nm, $\lambda_{em}$ = 500–650 nm). The relative emission responses ($\lambda_{ex}$ = 485 nm, $\lambda_{em}$ = 515 nm) from panels G–I were used to determine the apparent dissociation constants of nitrate binding to (J) ChlorON-1 ($K_d$ not determined, $R^2$ = 0.95), (K) ChlorON-2 ($K_d$ = 117 ± 18), and (L) ChlorON-3 ($K_d$ = 54 ± 4). The average of four technical replicates from two protein preparations with standard error of the mean is reported (Table S5).

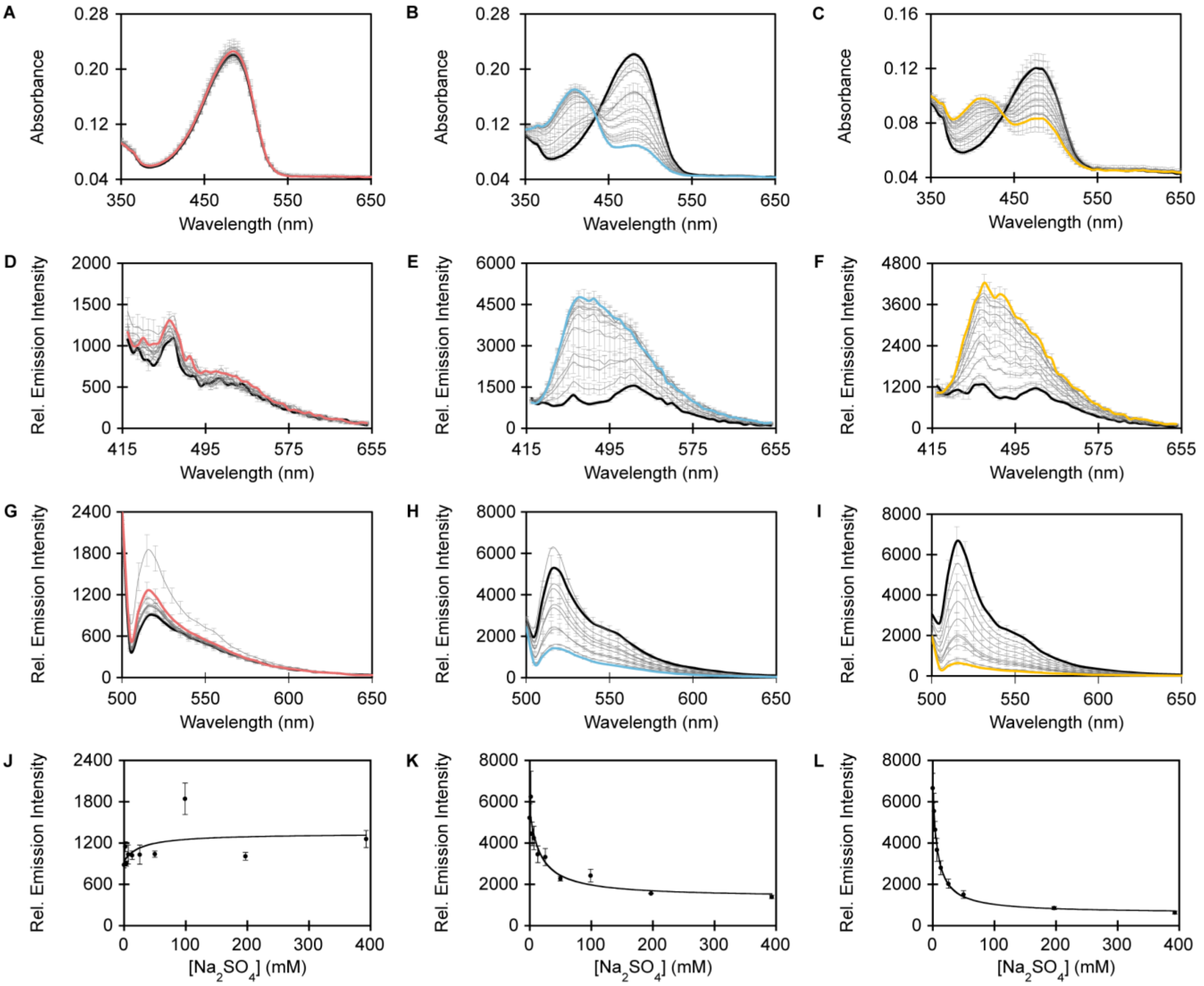


**Figure S19.** Spectroscopic characterization of ~10 µM ChlorON-1, ChlorON-2, and ChlorON-3 in the presence of 0 (bold black), 1.0, 2.9, 5.9, 12.3, 24.5, 49, 98, 196, and 392 mM (red, blue, and yellow, respectively) $Na_2SO_4$ in 50 mM sodium phosphate buffer at pH 7. Absorption spectra of (A) ChlorON-1, (B) ChlorON-2, and (C) ChlorON-3. Emission spectra of (D) ChlorON-1, (E) ChlorON-2, and (F) ChlorON-3 ($\lambda_{ex}$ = 400 nm, $\lambda_{em}$ = 415–650 nm). Emission spectra of (G) ChlorON-1, (H) ChlorON-2, and (I) ChlorON-3 ($\lambda_{ex}$ = 485 nm, $\lambda_{em}$ = 500–650 nm). The relative emission responses ($\lambda_{ex}$ = 485 nm, $\lambda_{em}$ = 515 nm) from panels G–I were used to determine the apparent dissociation constants of sulfate binding to (J) ChlorON-1 ($K_d$ not determined), (K) ChlorON-2 ($K_d$ = 17 ± 2), and (L) ChlorON-3 ($K_d$ = 7.5 ± 1). The average of four technical replicates from two protein preparations with standard error of the mean is reported (Table S5).

**Table S5.** Summary of the anion selectivity of ChlorON-1, ChlorON-2, and ChlorON-3 in 50 mM sodium phosphate buffer at pH 7. The average of four technical replicates from two protein preparations with standard error of the mean (SEM) is reported.

| | **ChlorON-1** | | **ChlorON-2** | | **ChlorON-3** | |
|---|---|---|---|---|---|---|
| **Anion** | **$F_f/F_i$ ± SEM** | **$K_d$ ± SEM (mM)** | **$F_f/F_i$ ± SEM** | **$K_d$ ± SEM (mM)** | **$F_f/F_i$ ± SEM** | **$K_d$ ± SEM (mM)** |
| Chloride | 45 ± 3 | 285 ± 59 | 27 ± 5 | 55 ± 5 | 20 ± 1 | 30 ± 1 |
| Bromide | 26 ± 3 | 204 ± 38 | 30 ± 2 | 43 ± 1 | 15 ± 2 | 32 ± 2 |
| Iodide | 2.6 ± 0.3 | — | 6.3 ± 0.3 | 86 ± 4 | 3.8 ± 0.8 | 48 ± 7 |
| Nitrate | 2.2 ± 0.5 | — | 9.1 ± 1 | 117 ± 18 | 7.9 ± 1 | 54 ± 4 |
| Sulfate | no response | — | 0.32 ± 0.09 | 17 ± 2 | 0.13 ± 0.01 | 7.5 ± 1 |
| Acetate | no response | — | no response | — | no response | — |
| Citrate | no response | — | no response | — | no response | — |
| Phosphate[a] | no response | — | no response | — | no response | — |

[a]*Determined in 50 mM HEPES buffer at pH 7.*

**A**

atg cat cat cac cat cac cat ggt acc gag ctc **gga tcc** ATG GTG TCC AAG GGC GAG GAG GAC AAT ATG
Met His His His His His His Gly Thr Glu Leu **Gly Ser** Met Val Ser Lys Gly Glu Glu Asp Asn Met

GCC TCT CTG CCA GCC ACC CAC GAG CTG CAC ATC TTC GGC TCT ATC AAC GGC GTG GAC TTT GAT ATG GTG
1 Ala Ser Leu Pro Ala Thr His Glu Leu His Ile Phe Gly Ser Ile Asn Gly Val Asp Phe Asp Met Val

GGA CAG GGA ACC GGA AAC CCA AAT GAC GGC TAC GAG GAG CTG AAT CTG AAG TCT ACA AAG GGC GAT CTG
24 Gly Gln Gly Thr Gly Asn Pro Asn Asp Gly Tyr Glu Glu Leu Asn Leu Lys Ser Thr Lys Gly Asp Leu

CAG TTC AGC CCT TGG ATT CTG GTG CCA CAC ATC GGC TAT GGC TTT CAC CAG TAT CTG CCC TAC CCT GAC
47 Gln Phe Ser Pro Trp Ile Leu Val Pro His Ile Gly Tyr Gly Phe His Gln Tyr Leu Pro Tyr Pro Asp

GGC ATG TCT CCT TTC CAG GCC GCC ATG GTG GAT GGC AGC GGC TAC CAG GTG CAC AGG ACA ATG CAG TTT
70 Gly Met Ser Pro Phe Gln Ala Ala Met Val Asp Gly Ser Gly Tyr Gln Val His Arg Thr Met Gln Phe

GAG GAC GGC GCC TCC CTG ACC GTG AAC TAC CGC TAT ACA TAC GAG GGC TCT CAC ATC AAG GGA GAG GCA
93 Glu Asp Gly Ala Ser Leu Thr Val Asn Tyr Arg Tyr Thr Tyr Glu Gly Ser His Ile Lys Gly Glu Ala

CAG GTG AAG GGA ACC GGA TTC CCA GCA GAT GGA CCC GTG ATG ACC AAC AGC CTG ACA GCA GCA GAC TGG
116 Gln Val Lys Gly Thr Gly Phe Pro Ala Asp Gly Pro Val Met Thr Asn Ser Leu Thr Ala Ala Asp Trp

TGC CGG TCC AAG **AAG** ACA TAT CCC AAT GAT AAG ACC ATC ATC AGC ACC TTC AAG TGG TCC TAT ACC ACA
139 Cys Arg Ser Lys **Lys** Thr Tyr Pro Asn Asp Lys Thr Ile Ile Ser Thr Phe Lys Trp Ser Tyr Thr Thr

GGC AAC GGC AAG CGG TAC AGA AGC ACC GCC CGG ACC ACA TAT ACA TTT GCC AAG CCC ATG GCC GCC AAC
162 Gly Asn Gly Lys Arg Tyr Arg Ser Thr Ala Arg Thr Thr Tyr Thr Phe Ala Lys Pro Met Ala Ala Asn

TAT CTG AAG AAT CAG CCT ATG TAC GTG TTC **AGG** AAG ACC GAG CTG AAG CAC TCC AAG ACA GAG CTG AAT
185 Tyr Leu Lys Asn Gln Pro Met Tyr Val Phe **Arg** Lys Thr Glu Leu Lys His Ser Lys Thr Glu Leu Asn

TTC AAG GAG TGG CAG AAG GCC TTT ACC GAC GTG ATG GGC ATG GAT GAG CTG TAC AAG tga **gaa ttc**
208 Phe Lys Glu Trp Gln Lys Ala Phe Thr Asp Val Met Gly Met Asp Glu Leu Tyr Lys * **Glu Phe**

**B**

| Mutations | ChlorON-1 | ChlorON-2 | ChlorON-3 |
|---|---|---|---|
| K143 | Trp (TGG) | Arg (CGG) | Arg (CGG) |
| R195 | Leu (CTG) | Ile (ATT) | Leu (CTG) |

**Figure S20.** (A) The nucleotide (top row) and amino acid (bottom row) sequences of the mNG construct used for transfecting the FRT-CFTR cells. The nucleotide sequence for mNG (green) was cloned into the pcDNA3.1(+)-N-6His vector (black) between the BamHI and EcoRI restriction sites (black and bold) with an N-terminal poly-histidine tag and a C-terminal stop codon (*). The K143 and R195 sites targeted for mutagenesis are bolded and highlighted in yellow. (B) The mutations at the K143 and R195 sites and corresponding codons used to generate the ChlorON-1, ChlorON-2, and ChlorON-3 constructs from the mNG construct in panel A.

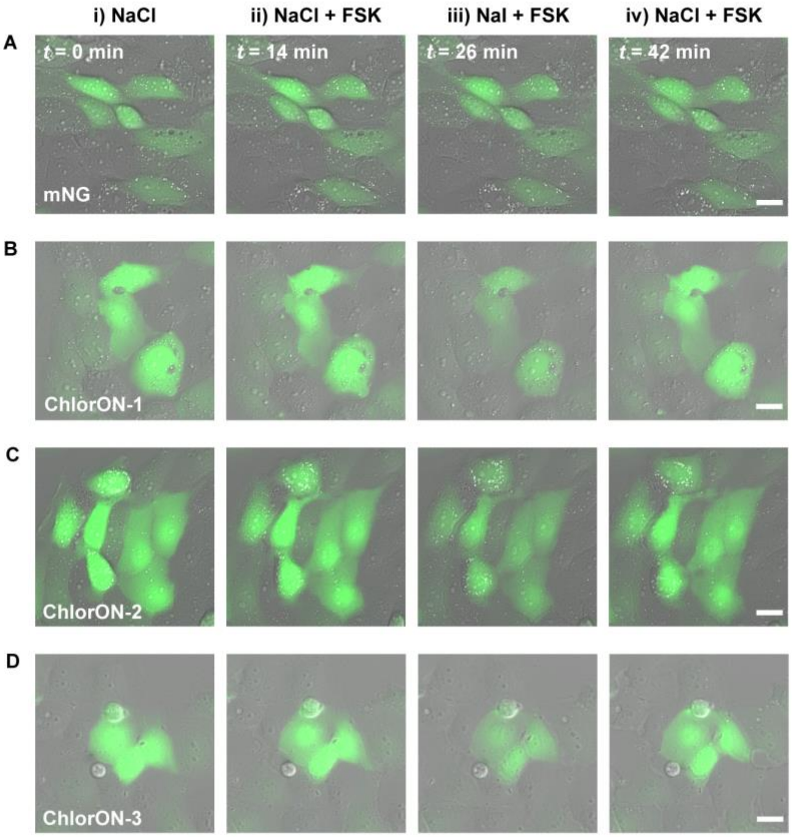


**Figure S21.** Representative fluorescence and differential interference contrast (DIC) images of live FRT-CFTR cells expressing (A) mNG, (B) ChlorON-1, (C) ChlorON-2, and (D) ChlorON-3 in a modified PBS buffer supplemented with (i) 137 mM NaCl at *t* = 0 min, (ii) 137 mM NaCl and 20 µM forskolin (FSK) at *t* = 14 min, (iii) 100 mM NaI, 37 mM NaCl, and 20 µM FSK at *t* = 26.5 min, and (iv) 137 mM NaCl and 20 µM FSK at *t* = 37.5 min. All experiments were carried out in a modified PBS buffer (2.7 mM KCl, 0.7 mM $CaCl_2$, 1.1 mM $MgCl_2$, 1.5 mM $KH_2PO_4$, 8.1 mM $Na_2HPO_4$, and 10 mM glucose) at pH 7.4 with the corresponding supplementation listed in each panel. Scale bar = 20 µm. Note: Exposure times are consistent within but variable between experiments (Figure 6 and Supplemental Video 1–4).

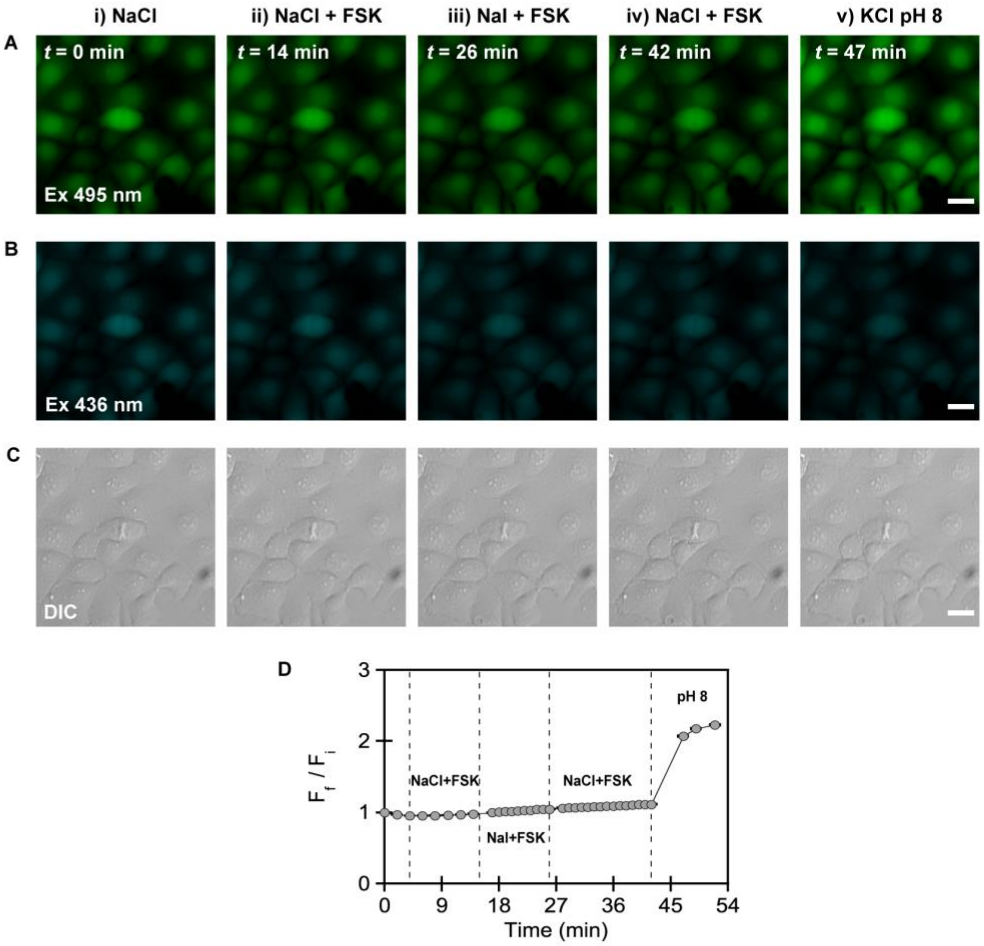


**Figure S22.** Representative fluorescence and DIC images of live FRT-CFTR cells stained with 5 µM BCECF. Fluorescence images are shown for the excitation provided at (A) 495 nm ($F_{Ex495}$) and (B) 436 nm ($F_{Ex436}$) with the corresponding (C) DIC images for cells treated with (i) 137 mM NaCl at *t* = 0 min ($F_i$), (ii) 137 mM NaCl and 20 µM forskolin (FSK) at *t* = 14 min, (iii) 100 mM NaI, 37 mM NaCl, and 20 µM FSK at *t* = 26 min, (iv) 137 mM NaCl and 20 µM FSK at *t* = 42 min, and (v) 137 mM KCl clamping buffer at pH 8 at *t* = 47 min. Steps (i–iv) were carried out in a modified PBS buffer (2.7 mM KCl, 0.7 mM $CaCl_2$, 1.1 mM $MgCl_2$, 1.5 mM $KH_2PO_4$, 8.1 mM $Na_2HPO_4$, and 10 mM glucose) at pH 7.4 with the corresponding supplementation listed in each panel. Step (v) was carried out in a clamping buffer at pH 8 (137 mM KCl, 2.7 mM NaCl, 0.7 mM $CaCl_2$, 1.1 mM $MgCl_2$, 1.5 mM $KH_2PO_4$, and 8.1 mM $Na_2HPO_4$) supplemented with 5 µM valinomycin and 5 µM nigericin. Scale bar = 20 µm. (D) Plot of the BCECF fluorescence response ($F_f/F_i$) for the $F_{Ex495}/F_{Ex436}$ emission of n = 1,810 regions of interest (ROIs). The average fluorescence response with standard error of the mean is reported for all ROIs from three biological replicates. The vertical dashed lines correspond to the transition between each condition.
